# Cluster effect through the oligomerisation of bioactive disaccharide AMOR on pollen tube capacitation in *Torenia fournieri*

**DOI:** 10.1101/2024.01.30.574946

**Authors:** Akane G Mizukami, Shuhei Kusano, Shinya Hagihara, Tetsuya Higashiyama

## Abstract

Arabinogalactan proteins (AGPs) are plant-specific glycoproteins involved in cellular mechanics and signal transduction. There has been major progress in understanding the structure, synthesis, and molecular functions of their carbohydrate chains; however, the mechanisms by which they function as signalling molecules remain unclear. Here, methyl-glucuronosyl arabinogalactan (AMOR; Me-GlcA-β(1,6)-Gal), a disaccharide structure at the end of AGP carbohydrate chains, was oligomerised via chemical synthesis. The biological activity of AMOR oligomers was enhanced via clustering of the carbohydrate chains. Furthermore, AMOR oligomers yielded a pollen tube morphology (i.e., callose plug formation) similar to that when cultured with native AMOR, suggesting it may be functionally similar to native AMOR.

## INTRODUCTION

The plant cell wall is not simply a structural component that provides mechanical support and protection to plant cells, it also contains cellular signalling molecules. Arabinogalactan proteins (AGPs), a subfamily of hydroxyproline-rich proteins highly glycosylated with type II arabinogalactans (AGs), are involved in various signalling pathways related to environmental responses, development, and reproduction in plants^1–5^. Type II AGs consist of a 1,3-β-galactan main chain with branches of 1,6-β-galactan side chains, which are further decorated with sugars such as arabinose, fucose, and glucuronic acid^5–9^. Type II AGs are synthesised as a sugar moiety of AGPs in the secretory pathway of plant cells by enzymes specific to each step of glycosylation^6,8^.

Many AGPs have been identified as signalling molecules^1,4,5^. For example, the signalling molecules involved in plant reproduction include tobacco transmitting-tissue specific proteins, 120-kDa glycoprotein, and PELPIII for pollen tube growth and pollen–pistil interactions^10–14^; *Arabidopsis thaliana* (At)AGP18 for female gametogenesis^15,16^; AtAGP6, AtAGP11, AtFLA3, rice MTR1, and AtAGP11 homolog *Brassica campestris* MF8 for pollen development^17–22^; AtENODL11/12/13/14/15 for pollen tube reception by the ovule^23^; and AtAGP4 (JAGGER) for polytubey block^24^. However, the functions of AGs in AGPs remain elusive. Knockout of enzymes involved in the initial steps of AG glycosylation show severe defects in plant development and reproduction, suggesting that type II AGs are critical for the bioactivity of AGPs^25–28^. Consistently, the distribution of AG structures visualised via immunostaining with monoclonal antibodies suggests tissue-specific accumulation of specific AG structures^3,28–30^. However, the functions of specific AG structures are largely unknown.

The heterogeneity of native polysaccharides, including their branch structures, impedes structural analyses. It is difficult to elucidate the structure and structure–activity relationships of individual native glycans. Research based on the chemical synthesis of sugar chains is important for closing this knowledge gap. Methyl-glucuronosyl arabinogalactan (AMOR; Me-GlcA-β(1,6)-Gal) is an AG derived from unfertilised female ovule tissue. In the flowering plant *Torenia fournieri*, AMOR enables the response of pollen tubes to cysteine-rich LURE attractant peptides^31,32^. A terminal disaccharide structure of AGs, 4-Me-GlcA-β(1,6)-Gal is necessary and sufficient for the bioactivity of AMOR. Native AMOR is likely a typical, high-molecular-weight AG of approximately 25 kDa based on a gel filtration study^31^, which potentially contains branched structures with multiple 4-Me-GlcA-β(1,6)-Gal termini. The activity of the disaccharide AMOR has not been compared with other synthetic polysaccharides with multiple 4-Me-GlcA-β(1,6)-Gal termini on a single molecule.

As indicated by the structure–activity relationship of synthetic AMOR, it is possible to introduce modifications to the second sugar, galactose, without impacting its activity^32^. However, the first sugar, methyl-glucuronic acid, cannot be modified without altering activity; for example, converting the terminal sugar to non-methylated glucuronic acid decreases the activity by approximately 1/1000, suggesting precise recognition by an unidentified receptor in pollen tubes^31,32^. The synthesis of AMOR derivatives with modifications at the second sugar would facilitate important aspects of AMOR research, such as the production of branched AMOR oligomers, visualisation of AMOR, and crosslinking with receptor candidates.

In this study, we successfully prepared AMOR derivatives with introduced azide groups and produced various AMOR oligomers, including dimers, trimers, and tetramers. When comparing the activity of branched oligomeric AMOR to that of unbranched disaccharide AMOR (monomer), the AMOR oligomers exhibited higher activities, although the total amount of 4-Me-GlcA-β(1,6)-Gal was same in each fraction. The AMOR oligomers showed novel activity, promoting callose plug formation in pollen tubes, which was also observed in ovule pre-culture medium containing native AMOR. Our results together suggest the synthetic AMOR oligomers are structurally and functionally similar to native AMOR.

## Results and discussion

### Chemical synthesis of AMOR oligomers

#### Design

Our previous structural–activity relationship analysis of AMOR revealed that the galactose portion of Me-GlcA-β(1,6)-Gal, which is the active site of AMOR, requires a pyranose backbone for its activity, whereas the 1-OH group is not essential for activity^32^. Therefore, we oligomerised AMOR through the 1-OH group of Gal. To secure an adequate supply of AMOR disaccharide for multimer formation, we achieved the stereoselective construction of the glycosidic linkage of Me-GlcA-β(1,6)-Gal. Rather than using a benzyl group, as in previous AMOR synthesis, we used a benzoyl group as the protecting group of the glycosyl donor (Me-GlcA), resulting in the selective production of β-anomer through the neighbouring group participation of the 2-O-benzoyl group. Inspired by click chemistry for multimer synthesis, we conceptualised copper-catalysed Huisgen cycloaddition, a component of click chemistry. We realised the efficient synthesis of AMOR multimers by reacting azide-tethered AMOR with bis-alkyne, tris-alkyne, and tetra-alkyne molecules.

### Synthesis of AMOR oligomers

To synthesise AMOR oligomers, we first synthesised azide-tethered AMOR with protecting groups (Scheme 1). We prepared the glycosyl donor **3** from tetra-acetyl-glucopyranosyl-bromide **1**. Koenigs– Knorr glycosylation of **1** and tetraethylene glycol monoazide with silver trifluoromethanesulfonate (AgOTf) afforded **2**. We transformed the protecting groups of **2** in four steps to synthesise the benzoyl-protected glycosyl acceptor **3**: the removal of acetyl groups; trityl protection of 6-OH; benzoyl protection; and the removal of the trityl group. Next, we synthesised the glycosyl donor **8** from benzylidene-protected glucopyranoside **4**. Three protection and deprotection steps yielded **5**: benzoyl protection; the removal of the benzylidene group; and trityl protection of 6-OH. The silver oxide-mediated methylation of **5** at 4-OH and the subsequent removal of the trityl group afforded **6**. Oxidation of the C-6 position and esterification of the resulting carboxylic group generated the glucuronic acid derivative **7**. The removal of the allyl group and trichloroacetimidation completed glycosyl donor synthesis. Glycosylation of **3** and **8** were performed using trimethylsilyl trifluoromethanesulfonate (TMSOTf) at −20°C in CH_2_Cl_2_, resulting in β-anomer selective formation of azide-tethered AMOR **9**.

**Scheme 1.**
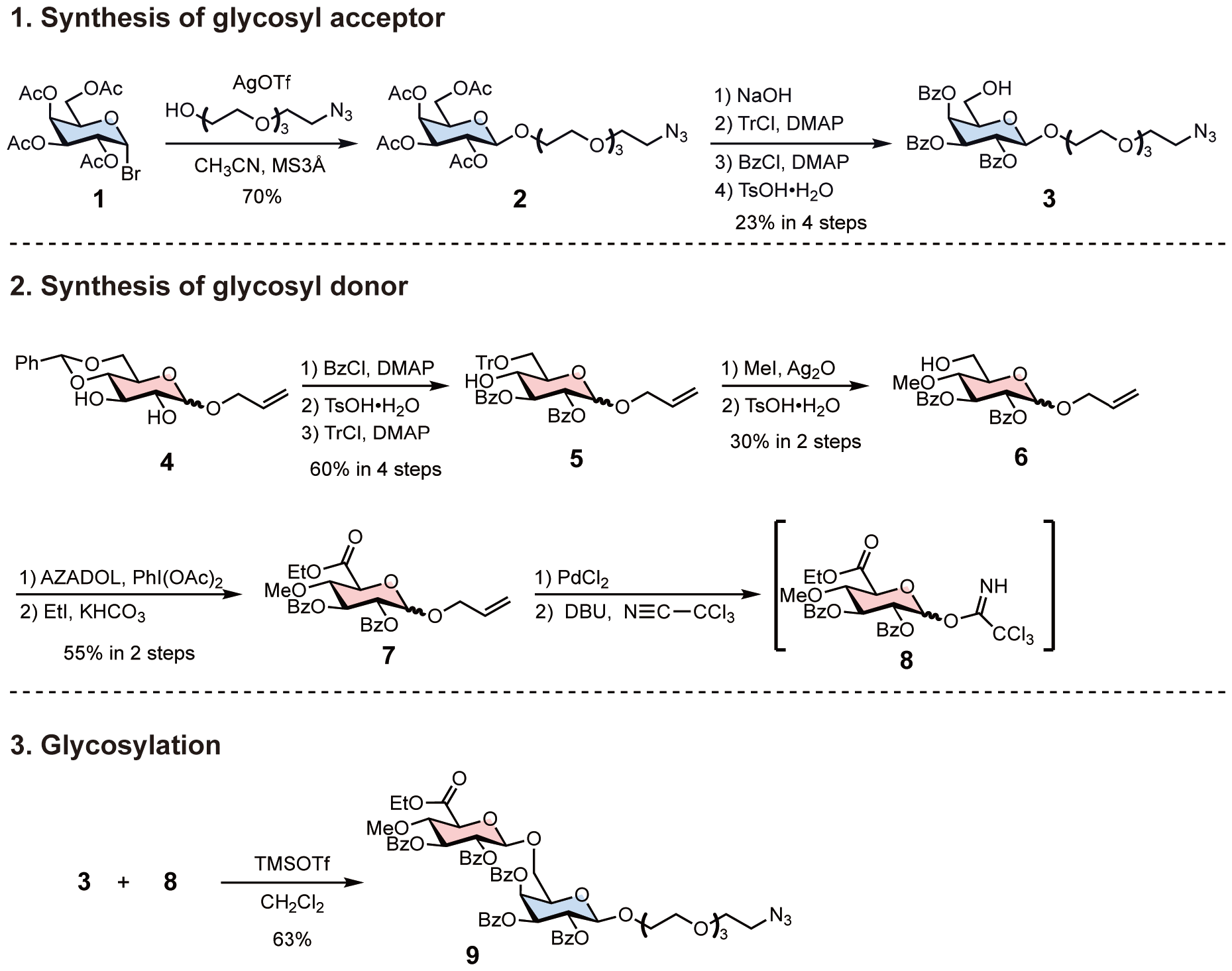
Synthesis of azide-tethered AMOR using protecting groups. Protecting the glycosyl donor with a benzoyl group increased the yield of β-anomer in the subsequent glycosylation reaction.

Subsequently, **9** was reacted with either bis-alkyne, tris-alkyne, or tetra-alkyne linkers to synthesise AMOR dimer (di-AMOR) **10**, AMOR trimer (tri-AMOR) **11**, and AMOR tetramer (tetra-AMOR) **12** with protecting groups, respectively (Scheme 2). The global deprotection of **10**–**12** was undertaken using LiOH•H_2_O to yield di-AMOR, tri-AMOR, and tetra-AMOR, respectively. The synthesised multivalent AMORs were characterised using ^1^H NMR and mass spectroscopy.

**Scheme 2.**
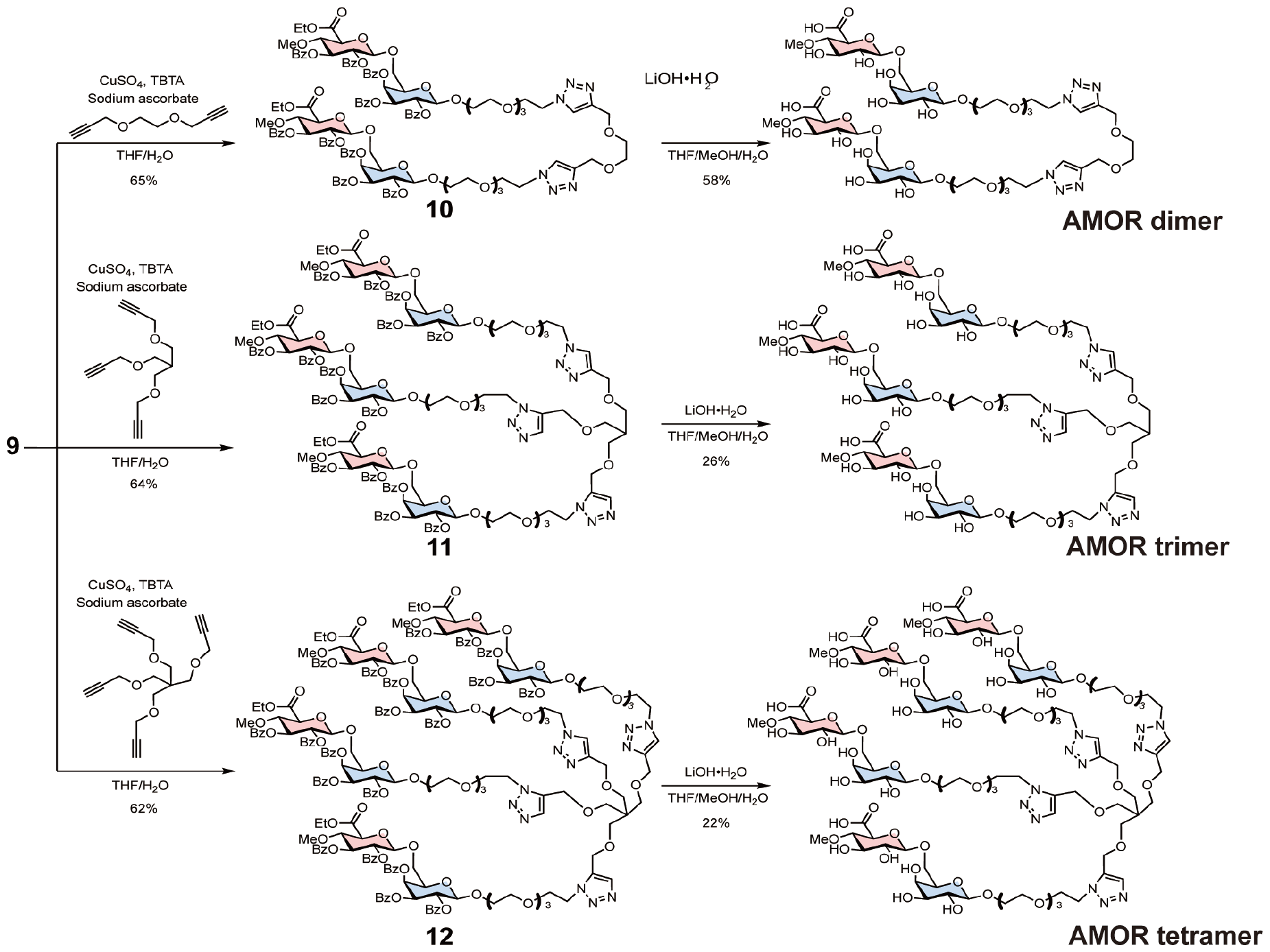
Synthesis of AMOR oligomers via the azide-alkyne click reaction.

### Comparison of AMOR monomer and oligomer activity

To investigate the impact of the cluster effect due to AMOR oligomerisation, we analysed the activity of AMOR monomer (mono-AMOR) and AMOR oligomers. To examine the response of pollen tubes, a freshly prepared ovule was placed using a glass needle near the tip of an elongating pollen tube growing through a cut style after overnight culture with chemically synthesised mono-AMOR or AMOR oligomers (Fig. 2A, B). Studies have shown that the AMOR activity of mono-AMOR saturates at 135 nM. Therefore, we measured AMOR activity at concentrations 10 and 100 times lower than that of mono-AMOR (Fig. 2C). Comparing equimolar concentrations of AMOR molecules, AMOR activity increased with the degree of oligomerisation. Thus, tetra-AMOR showed high AMOR activity at a low concentration of 3.38 nM, indicating that its structure readily clustered, leading to higher activity.

**Fig. 1.**
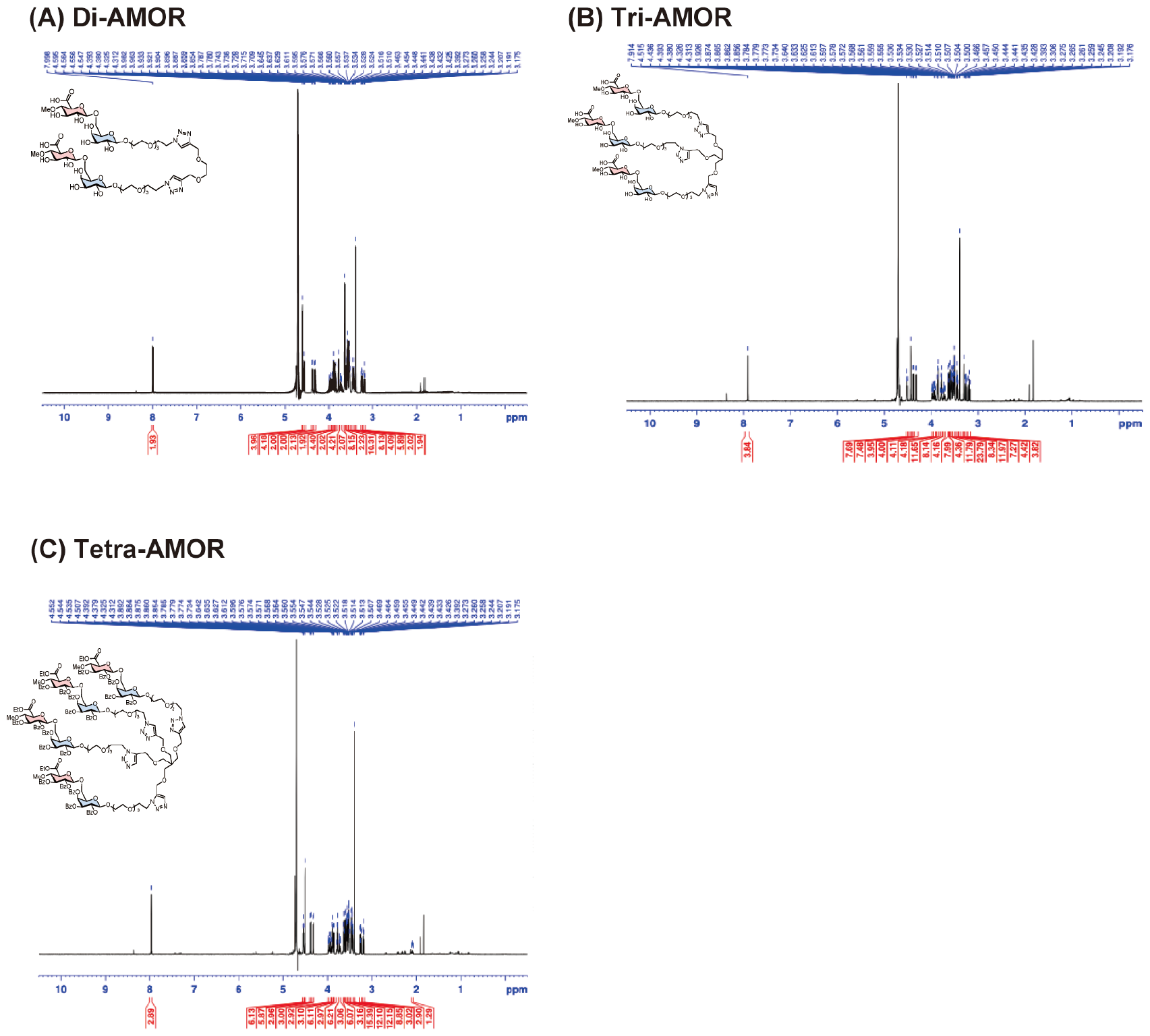
^1^H NMR (600 MHz, D_2_O) of **(A)** Di-AMOR, **(B)** Tri-AMOR, and **(C)** Tetra-AMOR.

**Fig. 2.**
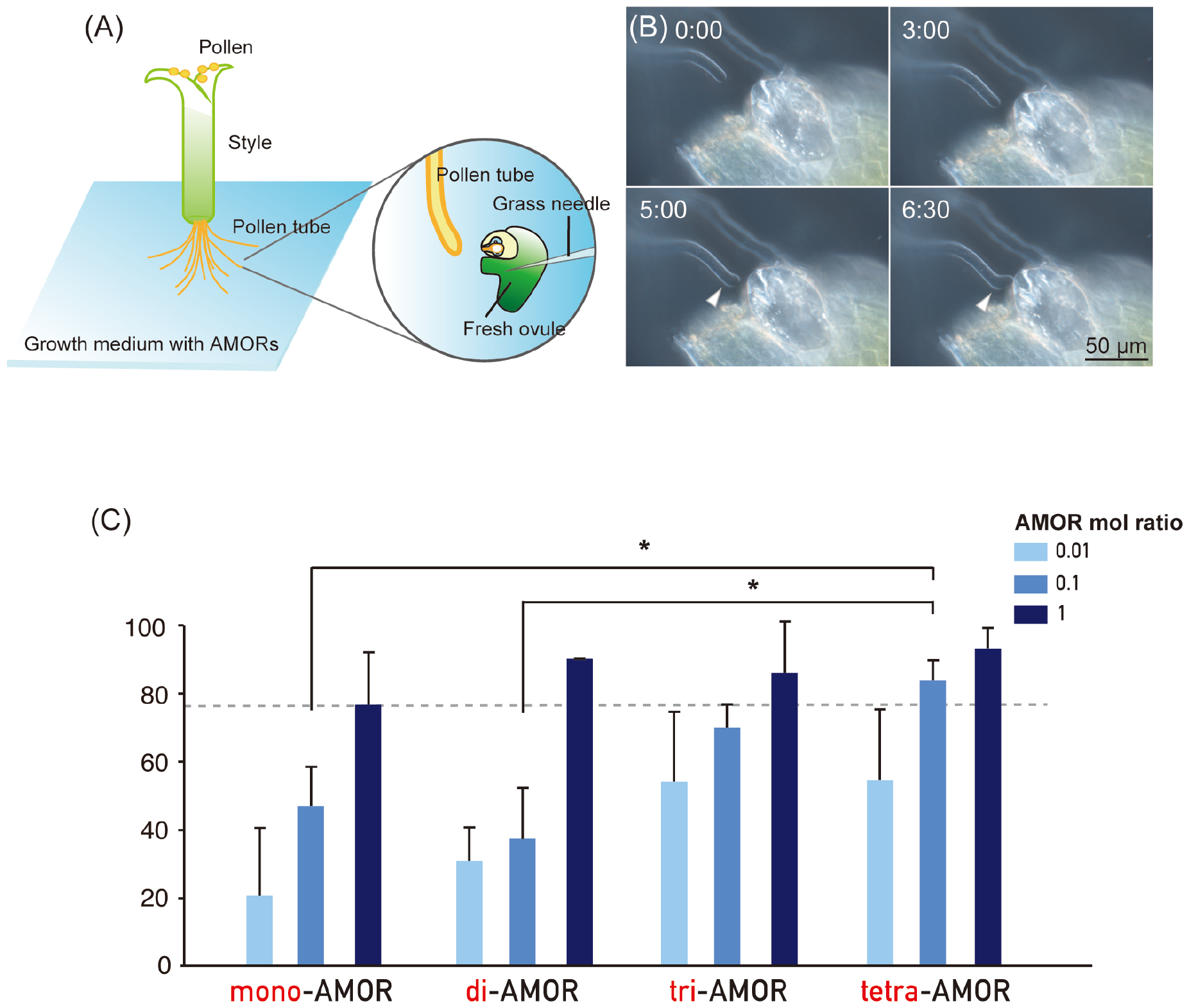
AMOR assay of AMOR monomer and AMOR oligomers. **(A)** Schematic representation of the AMOR assay for the quantification of pollen tube competency to ovular attraction. **(B)** Attraction of a competent pollen tube elongating in medium containing AMOR to a manipulated ovule. A manipulated ovule is placed in front of a pollen tube in the growth medium (0:00), which is allowed to elongate for a defined period (3:00), then change direction close to the micropyle (5:00), showing attraction to the tip of the micropyle (6:30). Time after placing an ovule is indicated as mm:ss. **(C)** AMOR activity of AMOR monomer and AMOR oligomers at concentrations of 1.35, 13.5, and 135 nM for mono-AMOR; 0.675, 6.75, and 67.5 nM for di-AMOR; 0.45, 4.5, and 45 nM for tri-AMOR; and 0.3375, 3.375, and 33.75 nM for tetra-AMOR. The concentration of mono-AMOR (disaccharide) of 135 nM was designed as a molar ratio of 1. In terms of the molar ratio of oligomer AMORs, the solutions contained the same total amounts of methyl-glucuronosyl galactose.Statistical analysis was performed using one-way analysis of variance with the *post hoc* Tukey’s HSD test and Bonferroni and Holm tests (*P < 0.05).

### Effect of AMORs on pollen tubes

We next conducted a morphological analysis to investigate the effects of AMORs on pollen tube growth. The distance between callose plugs in in vitro and semi-in vitro pollen tubes elongating in the growth medium was measured using aniline blue staining (Fig. 3A). Pollen tubes cultured in medium without AMORs had significantly shorter distances between callose plugs when passing through the style (Fig. 3B). In contrast, the presence or absence of AMORs had no effect on the distance between callose plugs in pollen tubes that did not traverse the style (Fig. 3B). These findings reveal that callose plug synthesis in pollen tubes is actively regulated by AMORs only when passing through the style. Interestingly, when using growth medium cultured with two-flowered ovules, even when the pollen tubes did not pass through the style (304.7 ± 121.6 µm), compared to cases in which they passed through the style (175.1 ± 99.8 µm), there was a significant decrease in the distance between the callose plugs of the pollen tubes (data not shown). These observations suggest that AMOR oligomers might functionally resemble native AMOR.

**Fig. 3.**
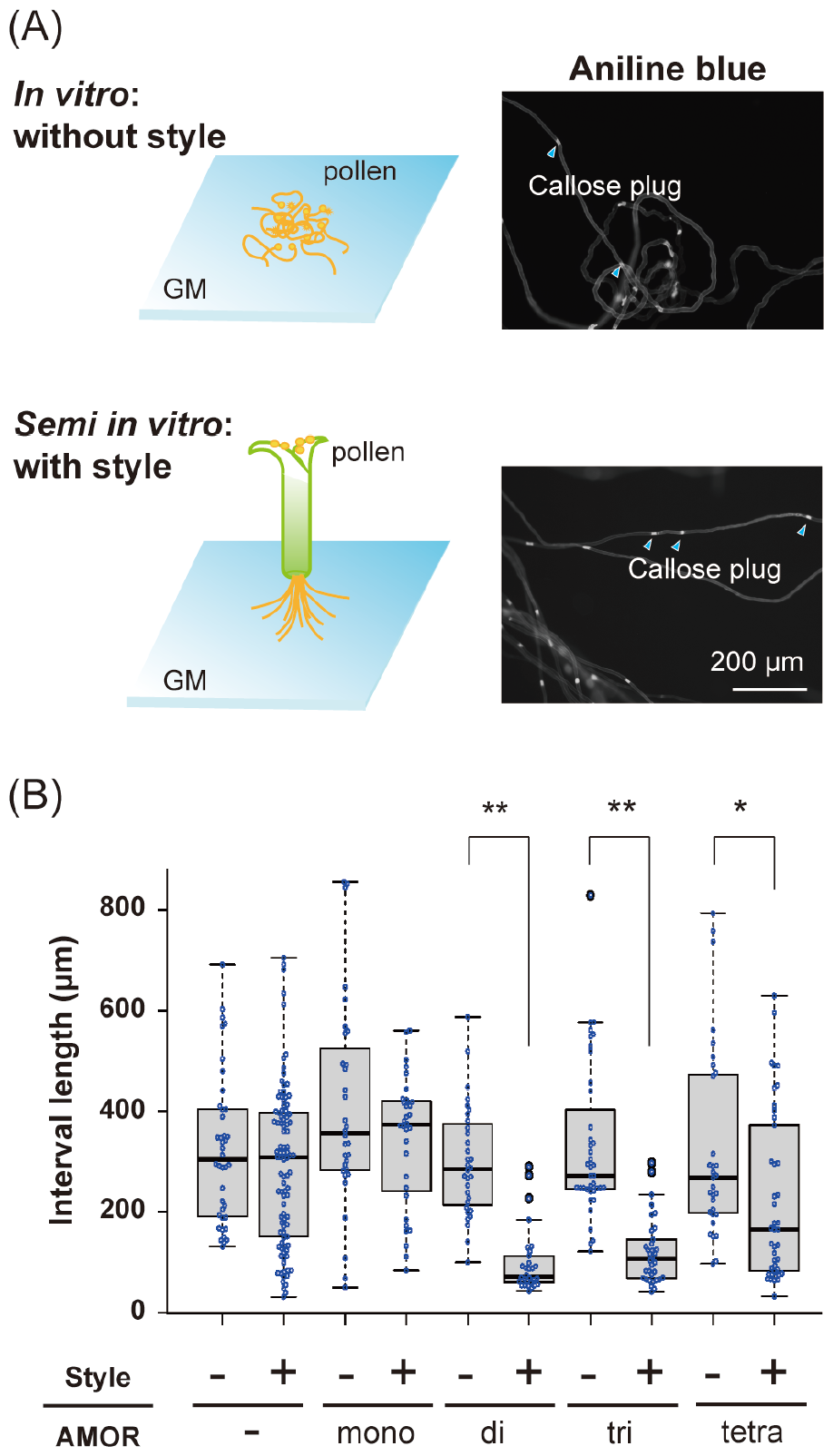
Distance between callose plugs in pollen tubes. **(A)** Schematic representation and aniline blue staining of *in vitro* and semi-*in vitro* pollen tubes. *In vitro* pollen tubes are directly germinated on growth medium (GM) containing AMOR and its oligomers. Semi-*in vitro* pollen tubes are generated by excising the pollinated pistil tip and placing it on GM containing AMOR and its oligomers, from which the pollen tubes elongate. Pollen tubes 16 h after the start of incubation were stained for callose plugs using aniline blue; three trials were made per condition, and the distance between callose plugs was measured for at least ten pollen tubes per trial. **(B)** Box-and whisker plots of distance between callose plugs. The concentrations of AMORs are 135 nM (mono-AMOR), 67.5 nM (di-AMOR), 45 nM (tri-AMOR), 33.75 nM (tetra-AMOR). Statistical test: Student’s t test (^*^p < 0.05, ^**^p < 0.01)

In general, glycan–protein interactions are weak. However, high affinity is achieved through the cluster glycoside effect, which amplifies the interaction by making the ligands and receptors multivalent^33–35^. The increased bioactivity of AMOR oligomers might represent an enhanced affinity of AMOR with unidentified receptor proteins or the activation of such AMOR receptors by their clustering, as previously suggested for some receptors such as the epidermal growth factor receptor in mammals^36,37^. Synthetic AMOR oligomers could provide novel approaches to better understand native AMOR and identify the AMOR receptor via chemical crosslinking of interacting proteins.

## Author contributions

A.G.M. and T.H. conceived of the idea; S.K. and S.H. synthesised and acquired the spectroscopic data of the AMOR and AMOR multimers; A.G.M. performed the AMOR assay and pollen tube observation.

## Acknowledgements

The authors thank Dr Kumi Matsuura (Riken, Japan), Dr Erika Toda (The University of Tokyo, Japan), Dr Daichi Susaki (Yokohama City University, Japan), and Dr Satohiro Okuda (The University of Tokyo, Japan) for their helpful discussions on this work. We also thank Ryo Yoshida, Akiko Takahashi, and Shoko Namiki for their help in maintaining the plants and other invaluable assistance. This work was supported by the Japan Society for the Promotion of Science KAKENHI (grant nos. 21K15119 to A.G.M., and 18H03997, 22H04980, and 22K21352 to T.H.) and the Japan Science and Technology Agency, Core Research for Evolutional Science and Technology (grant no. JPMJCR20E5 to T.H.).

