## Supplemental information for "Cluster effect through the oligomerisation of bioactive disaccharide AMOR on pollen tube capacitation in *Torenia fournieri*"

**Table of contents**

1. General S1
2. Synthetic scheme for AMOR oligomers S2-3
3. Synthesis procedures for AMOR oligomers S4-25
4. Bioassay S26
5. Reference S27

#### 1. General

Commercially available reagents were obtained from Tokyo Kasei, Wako Pure Chemical Industries Ltd., KANTO CHEMICAL CO., INC., Sigma-Aldrich Merck, and Nacalai tesque, and used without further purification.

The $^{1}$H and $^{13}$C{$^{1}$H} NMR were recorded on a Bruker AVANCE 600 (600 MHz for $^{1}$H, 150 MHz for $^{13}$C) spectrometer and a JEOL JNM-ECA500 (500 MHz for $^{1}$H, 125 MHz for $^{13}$C) spectrometer. Chemical shifts were reported in ppm (δ), and coupling constants were reported in Hz. $^{1}$H and $^{13}$C-resonances were referenced to solvent residual peaks for $CDCl_3$ ($^{1}$H, 7.26 ppm), $D_2O$ (1H, 4.70 Hz), and $CDCl_3$ ($^{13}$C, 77.16 ppm). Multiplicity and qualifier abbreviations are as follows: s = singlet, d = doublet, t = triplet, q = quartet, m = multiplet, doublet of doublets (dd), doublet of triplets (dt), doublet of doublet of doublets (ddd), doublet of doublet of doublet of doublets (dddd). Spectra were processed by Bruker Top-spin (Bruker) and Delta NMR software (JEOL).

High resolution mass analyses (HRMS) were submitted to the Mass Spectrometry Laboratory at RIKEN. For crude analysis, ultra high-performance liquid chromatography-mass spectrometry (UPLC/MS) was performed on a SHIMADZU LCMS-2020 equipped with a reverse phase C18 column (2.7 μm particle size, 2.1 x 100 mm) and a API/ESI mass spectrometry detector, and UV detector. MALDI-TOF MS was performed on a Bruker Daltonics UltrafleXtreme using α-cyano-4-hydroxycinnamic acid as a matrix.

Thin-layer chromatography was performed on Merck 60 F254 precoated silica gel plates. Reverse phase preparative Thin-layer chromatography was performed on Merck RP-8 modified silica gel plate coated with F254 indicator. Column chromatography was performed on open column using silica gel (Silica Gel 60 N; 63−210 mesh, KANTO CHEMICAL CO., INC. or 40−50 mesh, KANTO CHEMICAL CO., INC.).

### 2. Synthetic scheme for AMOR oligomers

#### *Scheme 1. Synthesis of azide-tethered AMOR with protection groups*

##### *Synthesis of glycosyl acceptor*

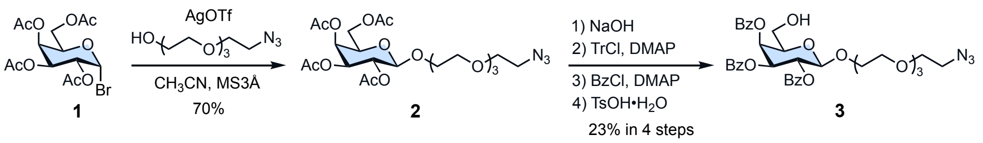


##### *Synthesis of glycosyl donor*

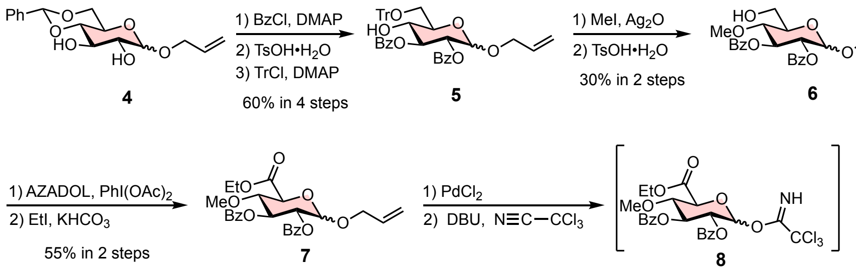


##### *Glycosylation*

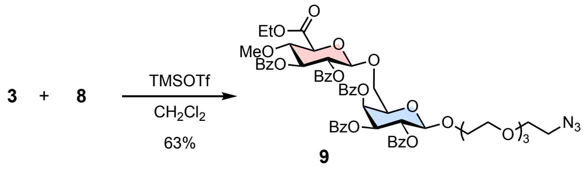

#### *Scheme 2. Synthesis of multivalent AMORs*

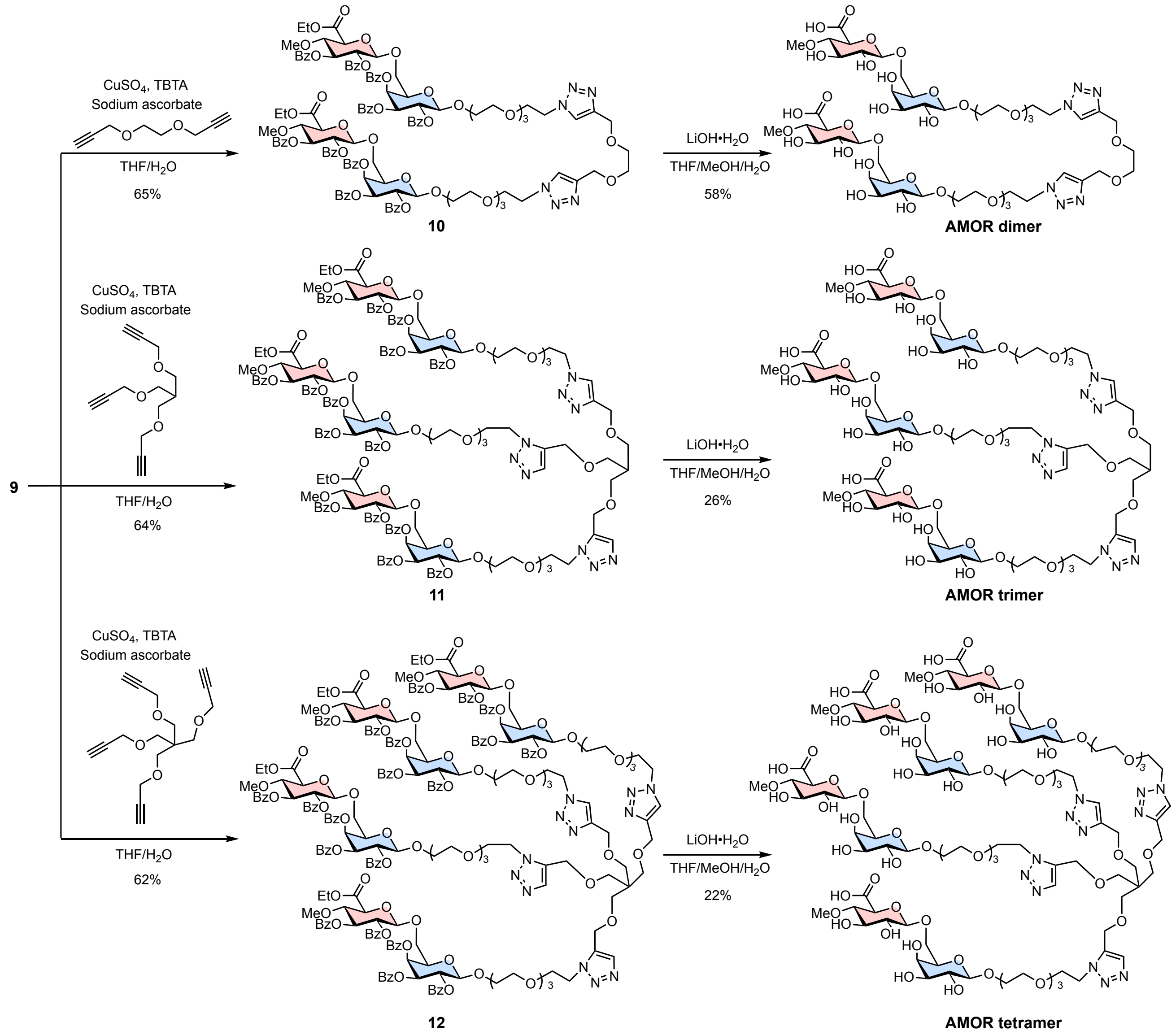

### 3. Synthesis procedures for AMOR oligomers

Compounds **4**, tetraethylene glycol mono-azide, and each alkyne linker were prepared as reported previously.[1–31–3]

**Synthesis of 2**

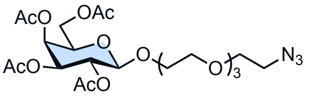


**2**

AgOTf (4.18 g, 16.3 mmol) was added to a mixture of **1** (6.1 g, 14.8 mmol) and tetraethylene glycol mono-azide (3.48 g, 19.8 mmol) in $CH_3CN$ (74 mL) and MS3Å at –20 ºC. This reaction mixture was stirred at –20 °C for 90 min and then allowed to slowly warm to room temperature. The reaction mixture was stirred for 18 h at room temperature, after which the mixture was filtrated through a pad of Celite® and concentrated. The residue was applied for column chromatography (silica gel, Hexane : AcOEt = 1 : 0 to 50 : 1 to 25 : 1) to afford **2** as a colorless oil (6.0 g, 70%); $^1$H NMR (500 MHz, $CDCl_3$) δ 5.38 (d, $J$ = 3.5 Hz, 1H), 5.21 (dd, $J$ = 8.0, 10.5 Hz, 1H), 5.01(dd, $J$ = 3.5, 10.5, 1H), 4.56 (d, $J$ = 8.0 Hz, 1H), 4.17 (dd, $J$ = 6.0, 10.5 Hz, 1H), 4.12 (d, $J$ = 7.0, 10.5 Hz, 1H), 3.95 (dt, $J$ = 4.0, 11.0 Hz, 1H), 3.91 (dd, $J$ = 6.0, 7.0 Hz, 1H), 3.77–3.72 (m, 1H), 3.69–3.62 (m, 12H), 3.39 (t, $J$ = 5.0 Hz, 2H), 2.15 (s, 3H), 2.06 (s, 3H), 2.05 (s, 3H), 1.98 (s, 3H); $^{13}$C NMR (125 MHz, $CDCl_3$) δ 170.5, 170.4, 170.3, 169.6, 101.5, 71.0, 70.8, 70.8, 70.4, 70.2, 69.2, 68.9, 67.2, 61.4, 50.8, 20.9, 20.8, 20.7; HRMS (ESI): calculated for $[M+Na]^+$ requires $m/z$ = 572.2068, found 572.2051.

**Synthesis of 3**

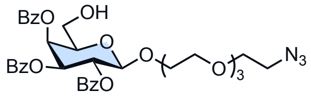


**3**

Step 1: Removal of acetyl group

To a solution of **2** (6.00 g, 10.9 mmol) in MeOH was added NaOH (1.96 g, 49.1 mmol) at room temperature. The mixture was stirred for 90 minutes at room temperature, after which the pH of the reaction mixture was neutralized with AcOH. The resulting mixture was concentrated *in vacuo* and the residue was applied for short column chromatography (silica gel, $CHCl_3$ : MeOH = 10 : 1 to 5 : 1). The roughly purified product (3.0 g) was use in next step without further purification.

Step 2: Trityl protection of 6-OH group

TrCl (6.58 g, 23.6 mmol) was added to a mixture of the above product (3.0 g, ~7.87 mmol) and DMAP (96.1 mg, 0.787 mmol) in pyridine (26 mL) at room temperature. The reaction mixture was stirred for 16 hours at room temperature, after which the reaction was quenched with an excess amount of MeOH and concentrated *in vacuo*. The residue was applied for short column chromatography (silica gel, $CHCl_3$ : MeOH = 10 : 1 to 5 : 1). The roughly purified product was use in next step without further purification.

Step 3: Benzoyl protection

BzCl (3.2 mL, 27.5 mmol) was slowly added to a mixture of the above product and DMAP (96.1 mg, 0.787 mmol) in pyridine (31 mL) at 0 ºC. This reaction mixture was stirred at 0 °C for 15 minutes and then warmed up to room temperature. After being stirred for 18 hours at room temperature, the reaction mixture was diluted with EtOAc and washed with water and brine. The organic layer was dried over $Na_2SO_4$, filtrated, and concentrated *in vacuo*. The residue was used in next step without any purification.

Step 4: Removal of trityl group

To a solution of the above product in $CH_2Cl_2$ (39 mL) and MeOH (39 mL) was added TsOH•$H_2O$ (271 mg, 1.57 mmol) at room temperature. After being stirred for 3 hours at room temperature, TsOH•$H_2O$ (700 mg, 4.07 mmol) was added to complete the reaction. The mixture was further stirred for 12 hours, after which the reaction mixture was diluted with $CH_2Cl_2$ and washed with sat. $NaHCO_3$ and brine. The organic layer was dried over $Na_2SO_4$, filtrated, and concentrated *in vacuo*. The residue was applied for column chromatography (silica gel, $CHCl_3$ : AcOEt = 2 : 1 to 1 : 1) to provide **3** as a white foam (1.74 g, 23% in four steps); $^1$H NMR (600 MHz, $CDCl_3$) δ 8.11 (d, $J$ = 7.5 Hz, 2H), 7.99 (d, $J$ = 7.5 Hz, 2H), 7.81 (d, $J$ = 7.5 Hz, 2H), 7.62 (t, $J$ = 7.5 Hz, 1H), 7.52 (t, $J$ = 7.5 Hz, 1H), 7.49 (dd, $J$ = 7.5, 7.5 Hz, 2H), 7.43 (t, $J$ = 7.5 Hz, 1H), 7.39 (dd, $J$ = 7.5, 7.5 Hz, 2H), 7.25 (dd, $J$ = 7.5, 7.5 Hz, 2H), 5.83 (dd, $J$ = 7.9, 10.2 Hz, 1H), 5.81 (d, $J$ = 3.4 Hz, 1H), 5.58 (dd, $J$ = 3.4, 10.2 Hz, 1H), 4.93 (d, $J$ = 7.9 Hz, 1H), 4.05–4.00 (m, 2H), 3.88–3.83 (m, 2H), 3.67–3.56 (m, 9 H), 3.52–3.45 (m, 4 H), 3.37 (t, $J$ = 4.8 Hz, 2H), 2.77 (t, $J$ = 6.8 Hz, 1H); $^{13}$C NMR (150 MHz, $CDCl_3$) δ 166.9, 165.7, 165.5, 134.0, 133.5, 133.4, 130.3, 129.9, 129.6, 128.9, 128.8, 128.8, 128.6, 128.5, 101.9, 74.2, 72.0, 70.8, 70.8, 70.7, 70.2, 69.6, 69.2, 60.8, 50.8; HRMS (ESI): calculated for $[M+Na]^+$ requires $m/z$ = 716.2431, found 716.2424.

**Synthesis of 5**

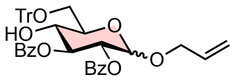


**5**

Step 1: Benzoyl protection

BzCl (4.7 mL, 16.3 mmol) was slowly added to a mixture of **4** (5.45 g, 17.7 mmol) and DMAP (216 mg, 1.77 mmol) in pyridine (89 mL) at 0 ºC. This reaction mixture was stirred at 0 °C for 15 minutes and then warmed up to room temperature. The reaction mixture was stirred for 18 hours at room temperature, after which the mixture was diluted with EtOAc and washed with water and brine. The organic layer was dried over $Na_2SO_4$, filtrated through a pad of silica gel, and concentrated *in vacuo*. The resulting product (6.52 g) was used in next step without further purification.

Step 2: Removal of benzylidene group

To a solution of the above product (6.52 g, ~12.6 mmol) in $CH_2Cl_2$ (67 mL) and MeOH (17 mL) was added TsOH•$H_2O$ (217 mg, 1.26 mmol) at room temperature. After being stirred for 3.5 hours at room temperature, TsOH•$H_2O$ (109 mg, 0.63 mmol) was added to complete the reaction. The mixture was further stirred for 1 hour, after which the reaction mixture was diluted with $CH_2Cl_2$ and washed with sat. $NaHCO_3$ and brine. The organic layer was dried over $Na_2SO_4$, filtrated, and concentrated *in vacuo*. The residue was used in next step without any purification.

Step 3: Trityl protection of 6-OH group

TrCl (5.27 g, 18.9 mmol) was added to a mixture of the above product and DMAP (154 mg, 1.26 mmol) in pyridine (42 mL) at room temperature. After being stirred for 12 hours at room temperature, TrCl (4.5 g, 16.1 mmol) was added to complete the reaction. The mixture was further stirred for 12 hours, after which the reaction was quenched with an excess amount of MeOH and concentrated *in vacuo*. The residue was applied for column chromatography (silica gel, Hexane: EtOAc = 6 : 1 to 3 : 1) to provide **5** as a white foam (7.08 g, 60% in three steps); $^{1}$H NMR (600 MHz, $CDCl_3$) δ 8.00–7.98 (m, 4H), 7.51–7.48 (m, 8H), 7.38–7.35 (m, 4H), 7.32 (dd, *J* = 7.4, 7.4 Hz, 6H), 7.25 (t, *J* = 7.4 Hz, 3H), 5.90–5.84 (m, 1H), 5.78 (dd, *J* = 9.0, 9.0 Hz, 1H), 5.31 (dddd, *J* = 1.3, 1.4, 1.5, 15.7 Hz, 1H), 5.27–5.25 (m, 2H), 5.16 (dddd, *J* = 1.3, 1.4, 1.5, 10.3 Hz, 1H), 4.25 (dddd, *J* = 1.5, 1.5, 5.1, 13.1 Hz, 1H), 4.05 (dddd, *J* = 1.3, 1.3, 6.0, 13.1 Hz, 1H), 3.96 (ddd, *J* = 4.0, 4.8, 9.1 Hz, 1H), 3.91 (ddd, *J* = 3.9, 9.1, 9.1 Hz, 1H), 3.49 (dd, *J* = 4.0, 10.1 Hz, 1H), 3.45 (dd, *J* = 4.8, 10.1 Hz, 1H), 2.83 (d, *J* = 3.9 Hz, 1H); $^{13}$C NMR (150 MHz, $CDCl_3$) δ 167.2, 166.1, 143.8, 133.7, 133.4, 133.4, 130.0, 130.0, 129.6, 129.4, 128.8, 128.5, 128.5, 128.1, 127.3, 117.7, 95.2, 87.3, 74.0, 71.6, 71.4, 70.5, 68.6, 64.0; HRMS (ESI): calculated for $[M+Na]^+$ requires *m/z* = 693.2464, found 693.2462.

**Synthesis of 6**

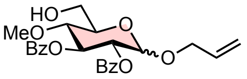


**6**

Step 1: Methylation

MeI (6.4 mL, 103 mmol) was added to a mixture of **5** (6.88 g, 10.3 mmol) and $Ag_2O$ (23.9 g, 103 mmol) in DMF (52 mL) at room temperature. After being stirred for 21 hours at room temperature, MeI (6.4 mL, 103 mmol) and $Ag_2O$ (23.9 g, 103 mmol) were added to complete the reaction. The mixture was further stirred for 18 hours, after which the mixture was filtrated through a pad of Celite®. The filtrate was diluted with EtOAc and washed with $H_2O$ in four times. The organic layer was dried over $Na_2SO_4$, filtrated through a pad of silica gel, and concentrated *in vacuo*. The residue was used in next step without further purification.

Step 2: Removal of benzylidene group

To a solution of the above product in $CH_2Cl_2$ (50 mL) and MeOH (50 mL) was added TsOH•$H_2O$ (710 mg, 2.06 mmol) at room temperature. The reaction mixture was stirred for 5 hours at room temperature, after which the mixture was diluted with $CH_2Cl_2$ and washed with sat. $NaHCO_3$ and brine. The organic layer was dried over $Na_2SO_4$, filtrated, and concentrated in vacuo. The residue was applied for column chromatography (silica gel, Hexane : AcOEt = 4 : 1 to 2 : 1) to provide **6** as a colorless oil (1.36 g, 30% in two steps); $^{1}$H NMR (600 MHz, $CDCl_3$) δ 8.02 (d, *J* = 7.7 Hz, 2H), 7.97 (d, *J* = 7.7 Hz, 2H), 7.52–7.49 (m, 2H), 7.39 (dd, *J* = 7.7, 7.7 Hz, 2H), 7.37 (dd, *J* = 7.7, 7.7 Hz, 2H), 5.99 (dd, *J* = 9.6, 9.6 Hz, 1H), 5.85–5.78 (m, 1H), 5.29–5.25 (m, 2H), 5.13 (d, *J* = 10.7 Hz, 1H), 5.11 (dd, *J* = 3.5, 10.3 Hz, 1H), 4.21 (dd, *J* = 6.8, 13.2 Hz, 1H), 4.02 (dd, *J* = 4.0, 13.2 Hz, 1H), 3.93–3.84 (m, 3H), 3.67 (dd, *J* = 9.6, 9.6 Hz, 1H), 3.47 (s, 3H), 1.89 (dd, *J* = 4.3, 7.9 Hz, 1H); $^{13}$C NMR (150 MHz, $CDCl_3$) δ 166.2, 165.8, 133.5, 133.4, 133.3, 130.0, 129.9, 129.8, 129.3, 128.6, 128.5, 117.8, 95.3, 77.9, 72.7, 72.3, 71.0, 68.8, 61.7, 60.7; HRMS (ESI): calculated for $[M+Na]^+$ requires *m/z* = 465.1525, found 465.1522.

**Synthesis of 7**

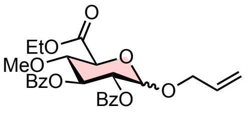

**7**

Step 1: Oxidation

AZADOL® (23.4 mg, 0.153 mmol) was added to a mixture of **6** (1.35 g, 3.05 mmol) and $PhI(OAc)_2$ (2.16 g, 6.71 mmol) in $CH_2Cl_2$ (8 mL) and 0.1 M phosphate buffer (pH 7.0, 8 mL) at 0 ºC. The reaction mixture was stirred for 2 hours at 0 ºC, after which the mixture was diluted with $CH_2Cl_2$ and washed with sat. $NaHCO_3$ and brine. The organic layer was dried over $Na_2SO_4$, filtrated, and concentrated *in vacuo*. The residue was used in next step without any purification.

Step 2: Esterification

Ethyl iodide (0.75 mL, 9.15 mmol) was added to a mixture of the above product and $KHCO_3$ (1.53 g, 51.3 mmol) in DMF (7.8 mL) at room temperature. The reaction mixture was stirred for 5 hours at room temperature, after which the mixture was diluted with EtOAc and washed with sat. $NaHCO_3$ and brine. The organic layer was dried over $Na_2SO_4$, filtrated, and concentrated *in vacuo*. The residue was applied for column chromatography (silica gel, Hexane : AcOEt = 4 : 1 to 2 : 1) to provide **7** as a colorless oil (820 mg, 55% in two steps); $^1H$ NMR (600 MHz, $CDCl_3$) δ 8.01 (dd, *J* = 1.0, 8.2 Hz, 2H), 7.97 (dd, *J* = 1.2, 8.2 Hz, 2H), 7.53–7.49 (m, 2H), 7.40 (dd, *J* = 8.2, 8.2 Hz, 2H), 7.37 (dd, *J* = 8.2, 8.2 Hz, 2H), 5.98 (dd, *J* = 9.7, 9.7 Hz, 1H), 5.85–5.79 (m, 1H), 5.32 (d, *J* = 3.7 Hz, 1H), 5.29 (dddd, *J* = 1.3, 1.4, 1.4, 17.1 Hz, 1H), 5.18 (dd, *J* = 3.5, 10.2 Hz, 1H), 5.15 (dddd, *J* = 1.3, 1.4, 1.4, 10.4 Hz, 1H), 4.35 (d, *J* = 9.7 Hz, 1H), 4.31 (q, *J* = 7.1 Hz, 2H), 4.27 (dddd, *J* = 1.4, 1.4, 5.0, 13.2 Hz, 1H), 4.05 (dddd, *J* = 1.3, 1.3, 6.2, 13.2 Hz, 1H), 3.85 (dd, *J* = 9.7, 9.7 Hz, 1H), 3.42 (s, 3H), 1.36 (t, *J* = 7.1 Hz, 3H); $^{13}C$ NMR (150 MHz, $CDCl_3$) δ 169.3, 166.0, 165.7, 133.5, 133.4, 133.2, 130.0, 129.9, 129.7, 129.2, 128.6, 128.6, 118.2, 95.7, 79.6, 72.1, 71.8, 70.3, 69.1, 62.0, 60.5, 14.3; HRMS (ESI): calculated for $[M+Na]^+$ requires *m/z* = 507.1631, found 507.1628.

**Synthesis of 8**

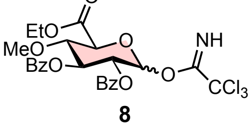

**8**

Step 1: Removal of allyl group

Under $N_2$ atmosphere, $PdCl_2$ (356 mg, 2.01 mmol) was added to a mixture of **7** (810 mg, 1.67 mmol) and NaOAc (342 g, 4.18 mmol) in AcOH (18 mL) and $H_2O$ (1.8 mL) at room temperature. The reaction mixture was stirred for 48 hours at room temperature, after which silica gel was added to the mixture. The volatile compounds were removed *in vacuo* and the resulting slurry of silica gel was applied for column chromatography (silica gel, Hexane : AcOEt = 6 : 1 to 3 : 1) to provide the deprotected product (718 mg). Although this product contained a small amount of impurity, it was used in next step without further purification.

Step 2: Trichloroacetimidation

DBU (24 µL, 0.162 mmol) was added to a solution of the above compound (718 mg, ~1.62 mmol) and $Cl_3CCN$ (1.6 mL, 16.2 mmol) in $CH_2Cl_2$ (16 mL) at 0 ºC. The reaction mixture was stirred at 0 °C for 15 minutes and then warmed up to room temperature. The reaction mixture was stirred for 4 hours at room temperature, after which the mixture was concentrated *in vacuo*. The residue was applied for short column chromatography (silica gel, Hexane : AcOEt = 4 : 1 to 2 : 1 + 1% $Et_3N$) to provide **7** as a white foam (234 mg); After the column purification, the resulting product was immediately used in next reaction due to the instability.

**Synthesis of 9**

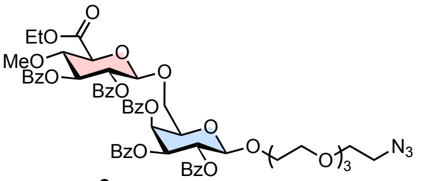

**9**

Under $N_2$ atmosphere, TMSOTf (15 µL, 16.3 mmol) was added to a mixture of **2** (520 mg, 0.750 mmol) and **8** (234 mg, 0.419 mmol) in $CH_2Cl_2$ (4 mL) at –20 ºC. This reaction mixture was stirred at –20 °C for 3 h and then allowed to slowly warm up to room temperature. The reaction mixture was stirred for 6 hours at room temperature, after which the mixture was quenched with an excess amount of $Et_3N$ and was then concentrated. The residue was applied for column chromatography (silica gel, Hexane : AcOEt = 4 : 1 to 2 : 1 to 1 : 1) to provide **9** as a colorless foam (294 mg, 63%); $^1$H NMR (600 MHz, $CDCl_3$) δ 8.05 (dd, *J* = 1.3, 8.3 Hz, 2H), 7.97 (dd, *J* = 1.3, 8.3 Hz, 2H), 7.94–7.92 (m, 4H), 7.74 (dd, *J* = 1.0, 8.2 Hz, 2H), 7.62–7.59 (m, 1H), 7.52–7.46 (m, 5H), 7.42–7.35 (m, 7H), 7.21 (t, *J* = 8.2 Hz, 2H), 5.80 (dd, *J* = 0.8, 3.4 Hz, 1H), 5.66 (dd, *J* = 7.9, 10.4 Hz, 1H), 5.60 (dd, *J* = 9.2, 9.2 Hz, 1H), 5.47 (dd, *J* = 3.4, 10.4 Hz, 1H), 5.38 (dd, *J* = 7.3, 9.2 Hz, 1H), 4.81 (d, *J* = 7.3 Hz, 1H), 4.72 (d, *J* = 7.9 Hz, 1H), 4.24–4.19 (m, 2H), 4.10 (ddd, *J* = 0.8, 3.4, 7.7 Hz, 1H), 4.05 (dd, *J* = 3.4, 10.9 Hz, 1H), 4.02 (d, *J* = 9.5 Hz, 1H), 3.91 (dd, *J* = 9.2, 9.5 Hz, 1H), 3.79 (dd, *J* = 7.7, 10.9 Hz, 1H), 3.65–3.63 (m, 3H), 3.59–3.58 (m, 2H), 3.51–3.45 (m, 3H), 3.40–3.30 (m, 11H), 1.27 (t, *J* = 7.1 Hz, 3H); $^{13}$C NMR (150 MHz, $CDCl_3$) δ 168.1, 165.7, 165.6, 165.6, 165.3, 165.2, 133.6, 133.5, 133.4, 133.3, 133.3, 130.2, 129.9, 129.9, 129.8, 129.8, 129.6, 129.3, 129.2, 129.0, 128.7, 128.6, 128.5, 128.5, 128.4, 101.4, 101.3, 79.2, 74.4, 74.3, 73.4, 72.0, 71.8, 70.7, 70.7, 70.6, 70.3, 70.1, 69.9, 69.4, 69.0, 68.8, 62.0, 60.6, 50.8, 14.2; HRMS (ESI): calculated for $[M+Na]^+$ requires *m/z* = 1142.3746, found 1142.3733.

**General procedure of click reaction for preparing 10–12**

Under $N_2$ atmosphere, sodium ascorbate (80 mol %) was added to a mixture of **9** (2.1–4.2 eq.), alkyne linker (1 eq.), $CuSO_4$ (20 mol %), and Tris[(1-benzyl-1H-1,2,3-triazol-4-yl)methyl]amine (TBTA, 40 mol %) in THF and $H_2O$ at room temperature. This reaction mixture was stirred at room temperature for 18 hours, after which the mixture was diluted with $CH_2Cl_2$ and washed with $H_2O$ and brine. The organic layer was dried over $Na_2SO_4$, filtrated, and concentrated *in vacuo*. The residue was applied for preparative TLC (silica gel, $CHCl_3$ : MeOH = 10 : 1) to afford **10–12**.

**Synthesis of 10**

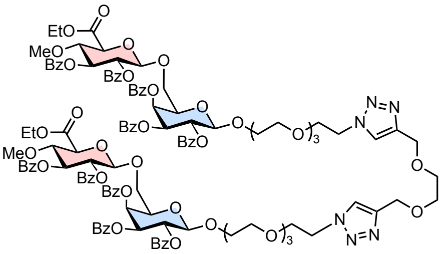

10

The reaction was performed with **9** (34.8 mg, 31.0 μmol), bis-alkyne linker (2 mg, 14.8 μmol), $CuSO_4$ (0.5 mg, 2.96 μmol), sodium ascorbate (2.3 mg, 11.8 μmol), TBTA (3.1 mg, 5.92 μmol) in THF (1 mL) and $H_2O$ (0.2 mL). After the purifications as described in the above, compound **10** was afforded as a colorless oil (23.0 mg, 65%); $^1$H NMR (600 MHz, $CDCl_3$) δ 8.04 (dd, *J* = 1.3, 8.3 Hz, 4H), 7.96 (dd, *J* = 1.3, 8.3 Hz, 4H), 7.93–7.90 (m, 8H), 7.73 (d, *J* = 1.1, 8.3 Hz, 4H), 7.70 (s, 2H), 7.61–7.58 (m, 2H), 7.52–7.44 (m, 10H), 7.41–7.32 (m, 14H), 7.21 (t, *J* = 8.3 Hz, 4H), 5.80 (dd, *J* = 0.7, 3.4 Hz, 2H), 5.66 (dd, *J* = 7.9, 10.4 Hz, 2H), 5.60 (dd, *J* = 9.2, 9.2 Hz, 2H), 5.47 (dd, *J* = 3.4, 10.4 Hz, 2H), 5.37 (dd, *J* = 7.3, 9.2 Hz, 2H), 4.81 (d, *J* = 7.3 Hz, 2H), 4.72 (d, *J* = 7.9 Hz, 2H), 4.65 (s, 4H), 4.49 (t, *J* = 5.4 Hz, 4H), 4.23–4.18 (m, 4H), 4.11 (dd, *J* = 3.4, 7.7 Hz, 2H), 4.04 (dd, *J* = 3.4, 10.9 Hz, 2H), 4.01 (d, *J* = 9.5 Hz, 2H), 3.90 (dd, *J* = 9.2, 9.5 Hz, 2H), 3.81 (t, *J* =5.4 Hz, 4H), 3.78 (dd, *J* = 7.7, 10.9 Hz, 2H), 3.68 (s, 4H), 3.66–3.63 (m, 2H), 3.52–3.50 (m, 4H), 3.48–3.42 (m, 6H), 3.40–3.32 (m, 14H), 3.28–3.26 (m, 4H), 1.26 (t, *J* = 7.1 Hz, 6H); $^{13}$C NMR (150 MHz, $CDCl_3$) δ 168.0. 165.6, 165.5, 165.3, 165.2, 144.9, 133.6, 133.5, 133.4, 133.3, 130.1, 129.9, 129.9, 129.8, 129.8, 129.6, 129.3, 129.3, 129.2, 128.8, 128.7, 128.6, 128.5, 128.5, 128.4, 123.9, 101.4, 101.3, 79.1, 74.5, 74.3, 73.3, 72.0, 71.8, 70.7, 70.6, 70.5, 70.4, 70.2, 69.9, 69.7, 69.5, 69.3, 69.0, 68.8, 64.7, 62.0, 60.6, 50.3, 14.2; HRMS (ESI): calculated for $[M+2H]^{2+}$ requires *m/z* = 1189.9283, found 1189.9282.

**Synthesis of 11**

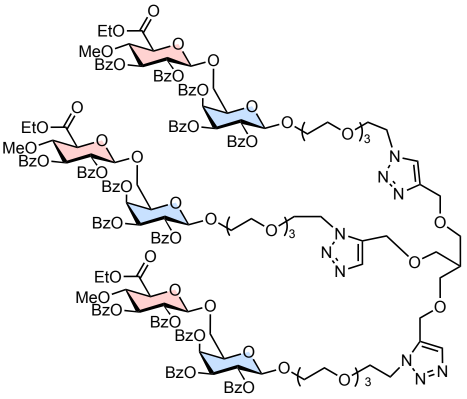

11

The reaction was performed with **9** (35.5 mg, 31.7 μmol), tris-alkyne linker (2.2 mg, 10.6 μmol), $CuSO_4$ (0.51 mg, 3.18 μmol), sodium ascorbate (2.5 mg, 12.7 μmol), TBTA (3.4 mg, 6.36 μmol) in THF (1 mL) and $H_2O$ (0.2 mL). After the purification as described in the above, compound **11** was afforded as a colorless oil (24.3 mg, 64%); $^1$H NMR (600 MHz, $CDCl_3$) δ 8.04 (dd, *J* = 1.3, 8.3 Hz, 6H), 7.96 (dd, *J* = 1.3, 8.3 Hz, 6H), 7.93–7.90 (m, 12H), 7.73 (dd, *J* = 1.1, 8.3 Hz, 6H), 7.67 (s, 3H), 7.61–7.58 (m, 3H), 7.52–7.44 (m, 15H), 7.41–7.32 (m, 21H), 7.21 (t, *J* = 8.3 Hz, 6H), 5.80 (dd, *J* = 0.8, 3.5 Hz, 3H), 5.66 (dd, *J* = 8.0, 10.4 Hz, 3H), 5.60 (dd, *J* = 9.2, 9.2 Hz, 3H), 5.47 (dd, *J* = 3.5, 10.4 Hz, 3H), 5.37 (dd, *J* = 7.3, 9.2 Hz, 3H), 4.81 (d, *J* = 7.3 Hz, 3H), 4.72 (d, *J* = 7.9 Hz, 3H), 4.54 (s, 6H), 4.49 (t, *J* = 5.3 Hz, 6H), 4.22–4.18 (m, 6H), 4.11 (dd, *J* = 3.4, 7.6 Hz, 3H), 4.04 (dd, *J* = 3.4, 11.0 Hz, 3H), 4.02 (d, *J* = 9.4 Hz, 3H), 3.90 (dd, *J* = 9.2, 9.4 Hz, 3H), 3.81 (t, *J* =5.3 Hz, 6H), 3.78 (dd, *J* = 7.6, 11.0 Hz, 3H), 3.66–3.63 (m, 3H), 3.54 (d, *J* = 5.9 Hz, 6H), 3.52–3.50 (m, 6H), 3.46–3.42 (m, 9H), 3.39–3.32 (m, 21H), 3.27–3.25 (m, 6H), 2.22–2.17 (m, 1H), 1.26 (t, *J* = 7.1 Hz, 9H); $^{13}$C NMR (150 MHz, $CDCl_3$) δ 168.1. 165.7, 165.6, 165.3, 165.2, 145.0, 133.7, 133.5, 133.4, 133.3, 130.2, 129.9, 129.9, 129.8, 129.8, 129.6, 129.3, 129.3, 129.2, 129.0, 128.7, 128.6, 128.6, 128.5, 128.4, 123.8, 101.4, 101.3, 79.2, 74.5, 74.3, 73.3, 72.0, 71.9, 70.7, 70.5, 70.5, 70.4, 70.2, 70.0, 69.5, 69.4, 69.0, 68.9, 68.8, 64.7, 62.0, 60.6, 50.3, 40.4, 14.2; HRMS (ESI): calculated for $[M+2H]^{2+}$ requires *m/z* = 1791.1433, found 1791.1451.

**Synthesis of 12**

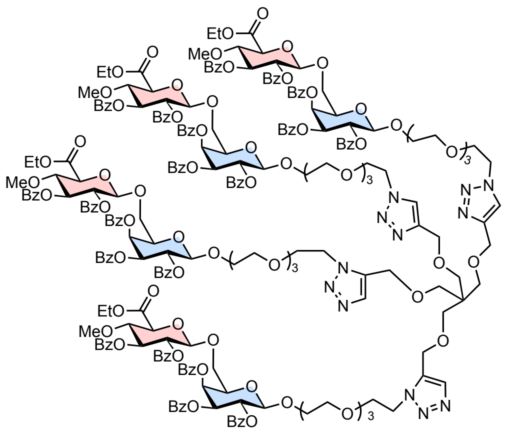

**12**

The reaction was performed with **9** (37.8 mg, 33.7 μmol), tetra-alkyne linker (2.3 mg, 8.03 μmol), $CuSO_4$ (0.51 mg, 3.21 μmol), sodium ascorbate (2.5 mg, 12.8 μmol), TBTA (3.4 mg, 6.42 μmol) in THF (2 mL) and $H_2O$ (0.4 mL). After the purification as described in the above, compound **12** was afforded as a colorless oil (23.8 mg, 62%); $^1$H NMR (600 MHz, $CDCl_3$) $^1$H NMR (600 MHz, $CDCl_3$) δ 8.04 (dd, *J* = 1.3, 8.4 Hz, 8H), 7.96 (dd, *J* = 1.3, 8.4 Hz, 8H), 7.93–7.89 (m, 16H), 7.73 (dd, *J* = 1.2, 8.4 Hz, 8H), 7.66 (s, 4H), 7.61–7.58 (m, 4H), 7.52–7.44 (m, 20H), 7.41–7.31 (m, 28H), 7.20 (t, *J* = 8.4 Hz, 8H), 5.80 (dd, *J* = 0.6, 3.6 Hz, 4H), 5.66 (dd, *J* = 8.0, 10.4 Hz, 4H), 5.60 (dd, *J* = 9.2, 9.2 Hz, 4H), 5.47 (dd, *J* = 3.5, 10.4 Hz, 4H), 5.37 (dd, *J* = 7.3, 9.2 Hz, 4H), 4.81 (d, *J* = 7.3 Hz, 4H), 4.72 (d, *J* = 7.9 Hz, 4H), 4.51 (s, 8H), 4.47 (t, *J* = 5.3 Hz, 8H), 4.22–4.18 (m, 8H), 4.11 (dd, *J* = 3.5, 7.7 Hz, 4H), 4.05 (dd, *J* = 3.5, 11.0 Hz, 4H), 4.01 (d, *J* = 9.5 Hz, 4H), 3.90 (dd, *J* = 9.2, 9.5 Hz, 4H), 3.81 (t, *J* =5.3 Hz, 8H), 3.78 (dd, *J* = 7.7, 11.0 Hz, 4H), 3.66–3.63 (m, 4H), 3.52–3.50 (m, 8H), 3.47–3.44 (m, 12H), 3.42–3.40 (m, 8H), 3.39–3.33 (m, 28H), 3.25–3.23 (m, 8H), 1.26 (t, *J* = 7.1 Hz, 12H); $^{13}$C NMR (150 MHz, $CDCl_3$) δ 168.0. 165.6, 165.6, 165.3, 165.2, 145.0, 133.7, 133.5, 133.4, 133.3, 130.2, 129.9, 129.9, 129.8, 129.8, 129.6, 129.3, 129.3, 129.2, 129.0, 128.7, 128.6, 128.6, 128.5, 128.4, 123.8, 101.4, 101.3, 79.2, 74.5, 74.3, 73.3, 72.0, 71.8, 70.7, 70.5, 70.5, 70.4, 70.2, 69.9, 69.5, 69.4, 69.0, 68.9, 68.8, 65.0, 62.0, 60.6, 50.2, 45.4, 14.2; calculated for $[M+3H]^{3+}$ requires *m/z* = 1590.5696, found 1590.5668.

**General procedure of hydrolysis reaction for preparing multivalent AMORs**

To a mixture of **10**–**12** (1 eq.) in THF, MeOH, and $H_2O$ was added LiOH•$H_2O$ (12–36 eq.). The reaction was monitored on LC-MS or MALDI-TOF MS. After the completion of the reaction, the mixture was quenched with AcOH and the pH was adjusted to around 4–5. The mixture was concentrated, and the residue was applied for preparative TLC (C8-modified silica gel, 5% $CH_3CN/H_2O$) and subsequent column chromatography (Sephadex$^{TM}$ LH-20, MeOH) to afford a multivalent AMORs. *The characterization of these compounds was performed with $^1$H NMR and mass spectroscopy.

**AMOR dimer**

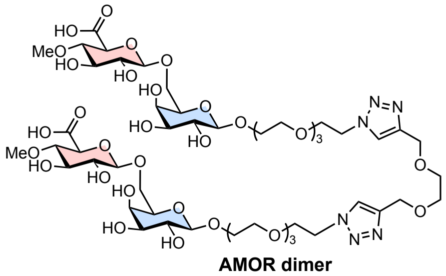

**AMOR dimer**

The reaction was performed with **10** (20.0 mg, 8.90 μmol) and LiOH•$H_2O$ (4.4 mg, 105 μmol) in THF (0.2 mL), MeOH (0.2 mL), and $H_2O$ (0.2 mL). After the purification as described in the above, **AMOR dimer** was afforded as a colorless oil (6.2 mg, 58%); $^1$H NMR (600 MHz, $D_2O$) δ 8.00 (s, 2H), 4.59 (s, 4H), 4.56 (t, *J* = 5.1 Hz, 4H), 4.38 (d, *J* = 7.9 Hz, 2H), 4.32 (d, *J* = 7.9 Hz, 2H), 3.97 (ddd, *J* = 4.1, 4.1, 11.7 Hz, 2H), 3.95–3.91 (m, 2H), 3.89 (t, *J* = 5.1 Hz, 4H), 3.85 (d, *J* = 3.4 Hz, 2H), 3.80–3.76 (m, 4H), 3.74–3.71 (m, 2H), 3.65–3.63 (m, 8H), 3.60 (d, *J* = 9.8 Hz, 2H), 3.58–3.55 (m, 10H), 3.54–3.51 (m, 8H), 3.46–3.42

(m, 4H), 3.39 (s, 6H), 3.26 (dd, *J* = 8.0, 9.2 Hz, 2H), 3.19 (dd, *J* = 9.5, 9.5 Hz, 2H); MS (ESI): calculated for $[M+H]^+$ requires *m/z* = 1281.5~~2~~, found ~~1281~~1280.7.

**AMOR trimer**

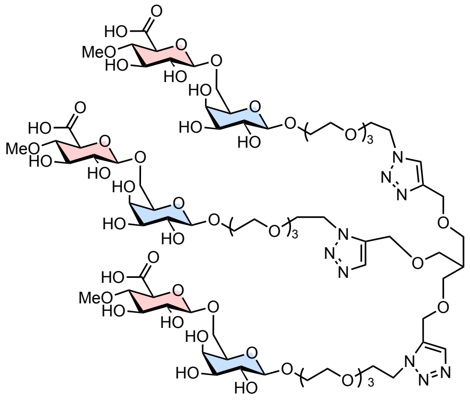


AMOR trimer

The reaction was performed with **11** (24.3 mg, 6.78 µmol) and LiOH• $H_2O$ (5.1 mg, 122 µmol) in THF (0.3 mL), MeOH (0.3 mL), and $H_2O$ (0.3 mL). After being stirred for 6 hours, LiOH• $H_2O$ (3.0 mg, 71.4 µmol) was added to complete the reaction. The mixture was further stirred for 24 hours. After the purification as described in the above, **AMOR trimer** was afforded as a colorless oil (3.4 mg, 26%); $^1H$ NMR (600 MHz, $D_2O$) δ 7.96 (s, 3H), 4.54 (t, *J* = 5.2 Hz, 6H), 4.50 (s, 6H), 4.38 (d, *J* = 7.9 Hz, 3H), 4.31 (d, *J* = 7.9 Hz, 3H), 3.97 (dt, *J* = 4.2, 11.7 Hz, 3H), 3.95–3.91 (m, 3H), 3.88 (t, *J* = 5.1 Hz, 6H), 3.86 (d, *J* = 3.5 Hz, 3H), 3.80–3.76 (m, 6H), 3.74–3.71 (m, 3H), 3.64 (t, *J* = 4.1 Hz, 6H), 3.60 (d, *J* = 9.8 Hz, 3H), 3.58–3.54 (m, 15H), 3.53–3.51 (m, 12H), 3.46–3.42 (m, 12H), 3.39 (s, 9H), 3.26 (dd, *J* = 7.9, 9.4 Hz, 3H), 3.19 (dd, *J* = 9.4, 9.4 Hz, 3H), 2.11–2.07 (m, 1H); MS (MALDI): calculated for $[M+H]^+$ requires *m/z* = 1934.8, found 1935.5.

**AMOR tetramer**

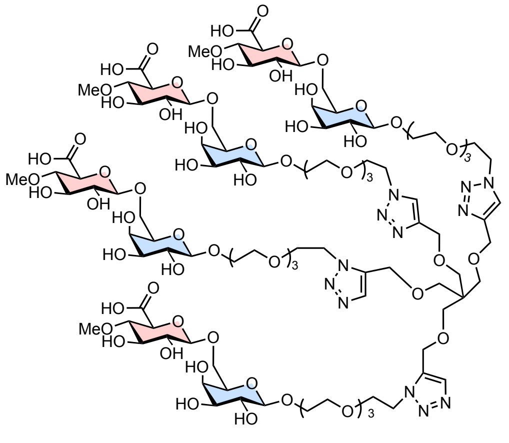


AMOR tetramer

The reaction was performed with **12** (23.8 mg, 4.99 µmol) and LiOH• $H_2O$ (5.1 mg, 122 µmol) in THF (0.3 mL), MeOH (0.3 mL), and $H_2O$ (0.3 mL). After being stirred for 6 hours, additional LiOH•$H_2O$ (6.0 mg, 143 µmol) was added to complete the reaction. The mixture was further stirred for 48 hours. After the purification as described in the above, **AMOR tetramer** was afforded as a colorless oil (2.8 mg, 22%); $^1H$ NMR (600 MHz, $D_2O$) δ 7.91 (s, 4H), 4.51 (t, *J* = 5.2 Hz, 8H), 4.44 (s, 8H), 4.38 (d, *J* = 7.9 Hz, 4H), 4.32 (d, *J* = 7.9 Hz, 4H), 3.97 (dt, *J* = 4.2, 11.8 Hz, 4H), 3.95–3.91 (m, 4H), 3.87–3.86 (m, 12H), 3.79–3.76 (m, 8H), 3.74–3.71 (m, 4H), 3.63 (d, *J* = 4.2 Hz, 8H), 3.60 (d, *J* = 9.8 Hz, 4H), 3.58–3.55 (m, 12H), 3.54–3.50 (m, 24H), 3.45–3.43 (m, 8H), 3.39 (s, 12H), 3.30 (s, 8H), 3.26 (dd, *J* = 7.9, 9.4 Hz, 4H), 3.19 (dd, *J* = 9.4, 9.4 Hz, 4H); MS (MALDI): calculated for $[M+H]^+$ requires *m/z* = 2574.0, found 2574.~~9~~6.

**2**: $^1$H NMR (500 MHz, $CDCl_3$)

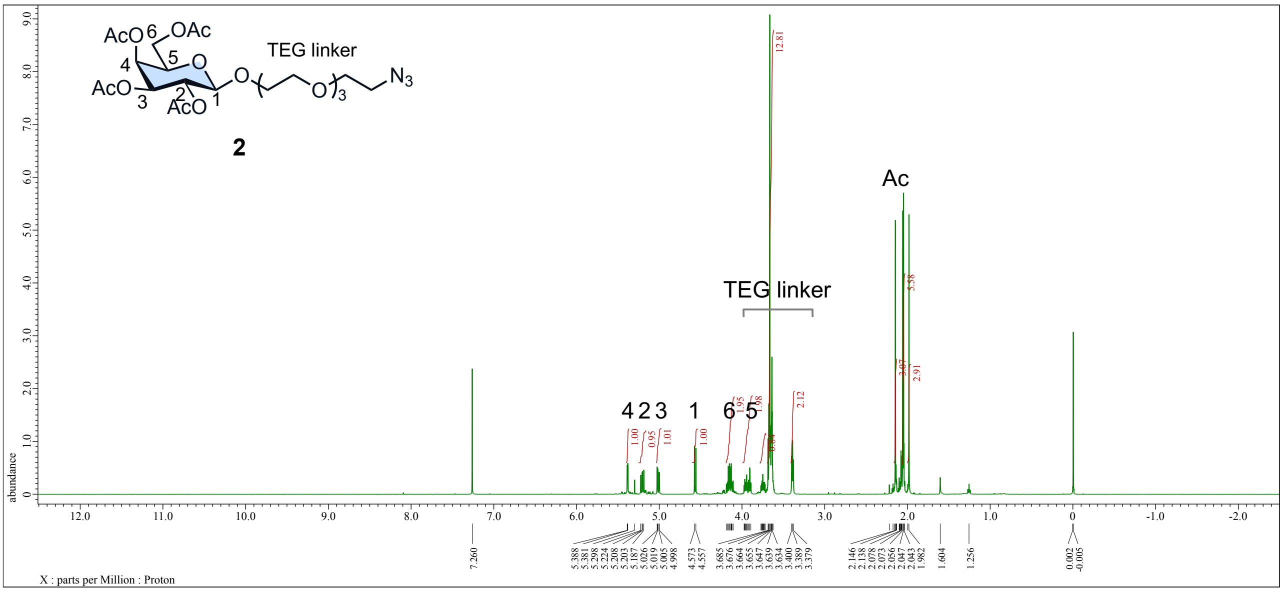


**2**: $^{13}$C NMR (125 MHz, $CDCl_3$)

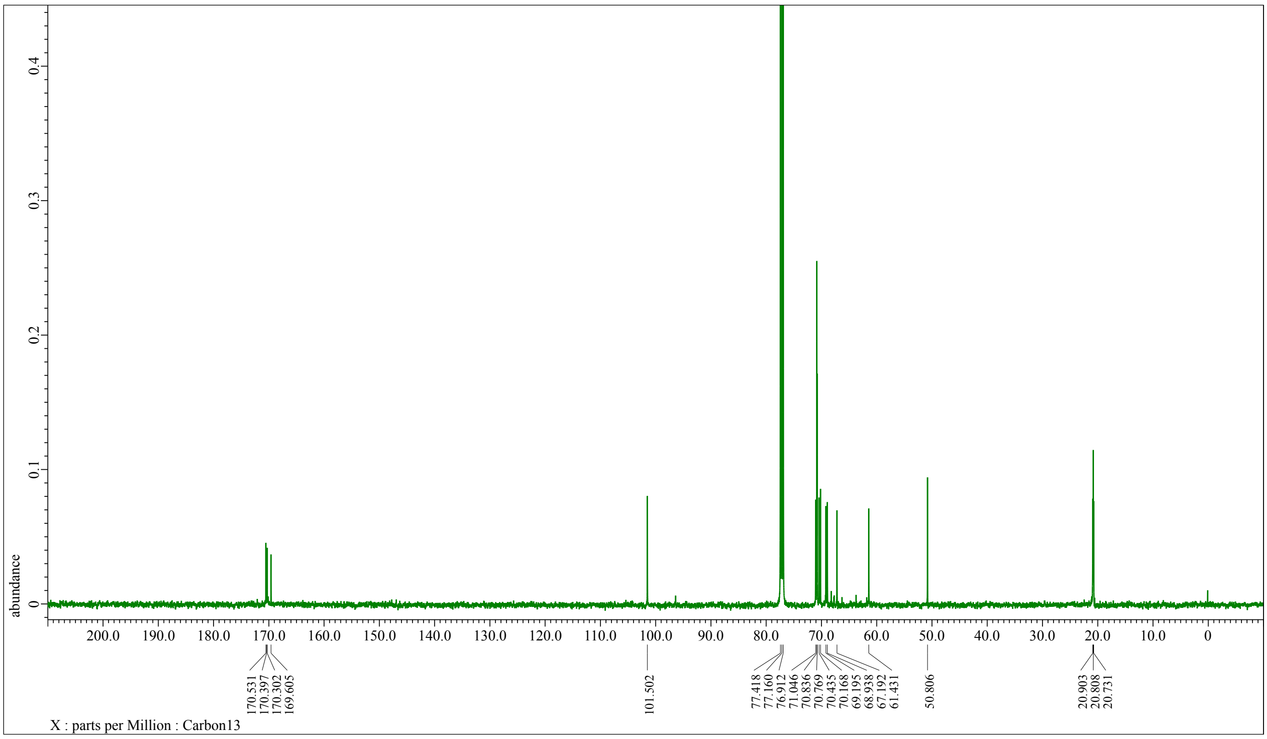

**3**: $^{1}$H NMR (600 MHz, CDCl$_3$)

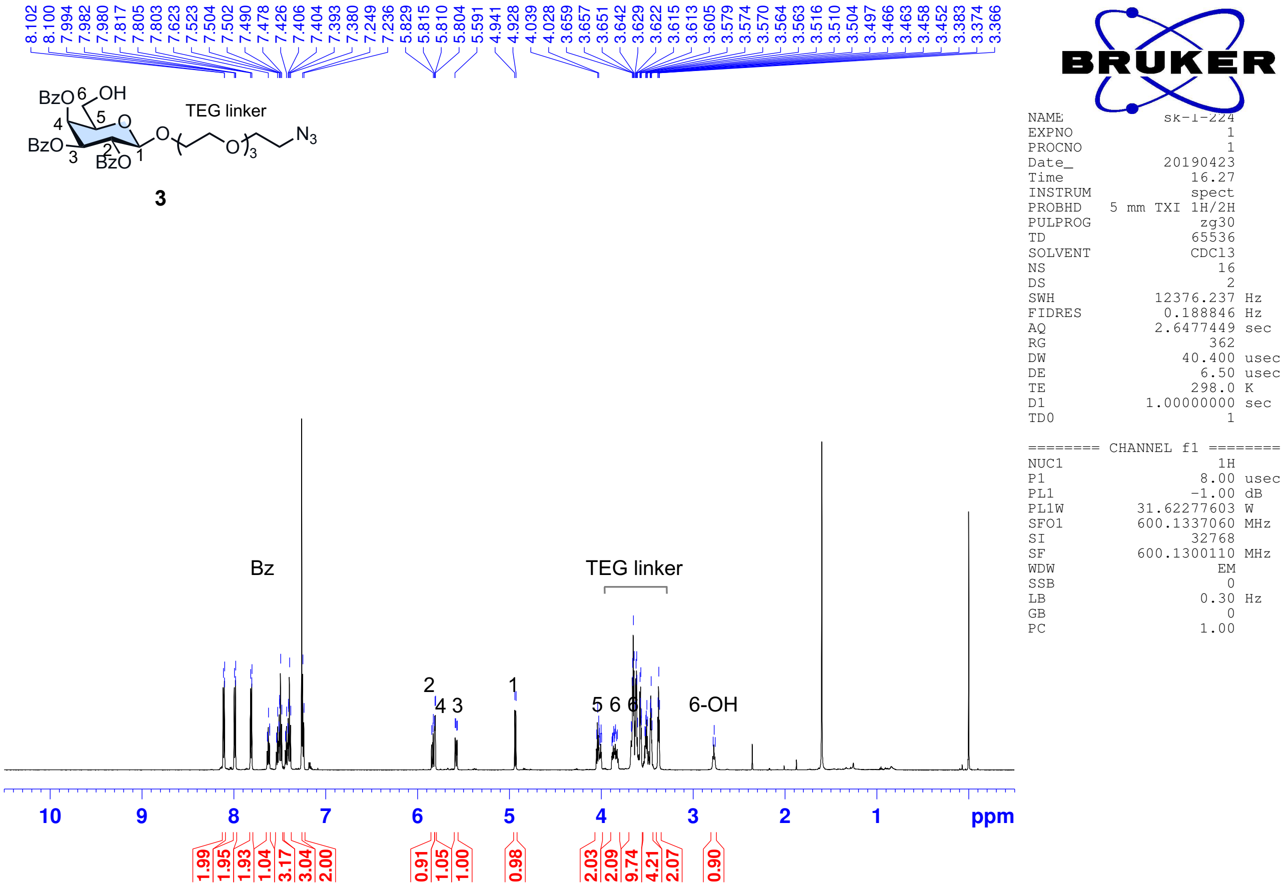


**3**: $^{13}$C NMR (150 MHz, CDCl$_3$)

1D 13C

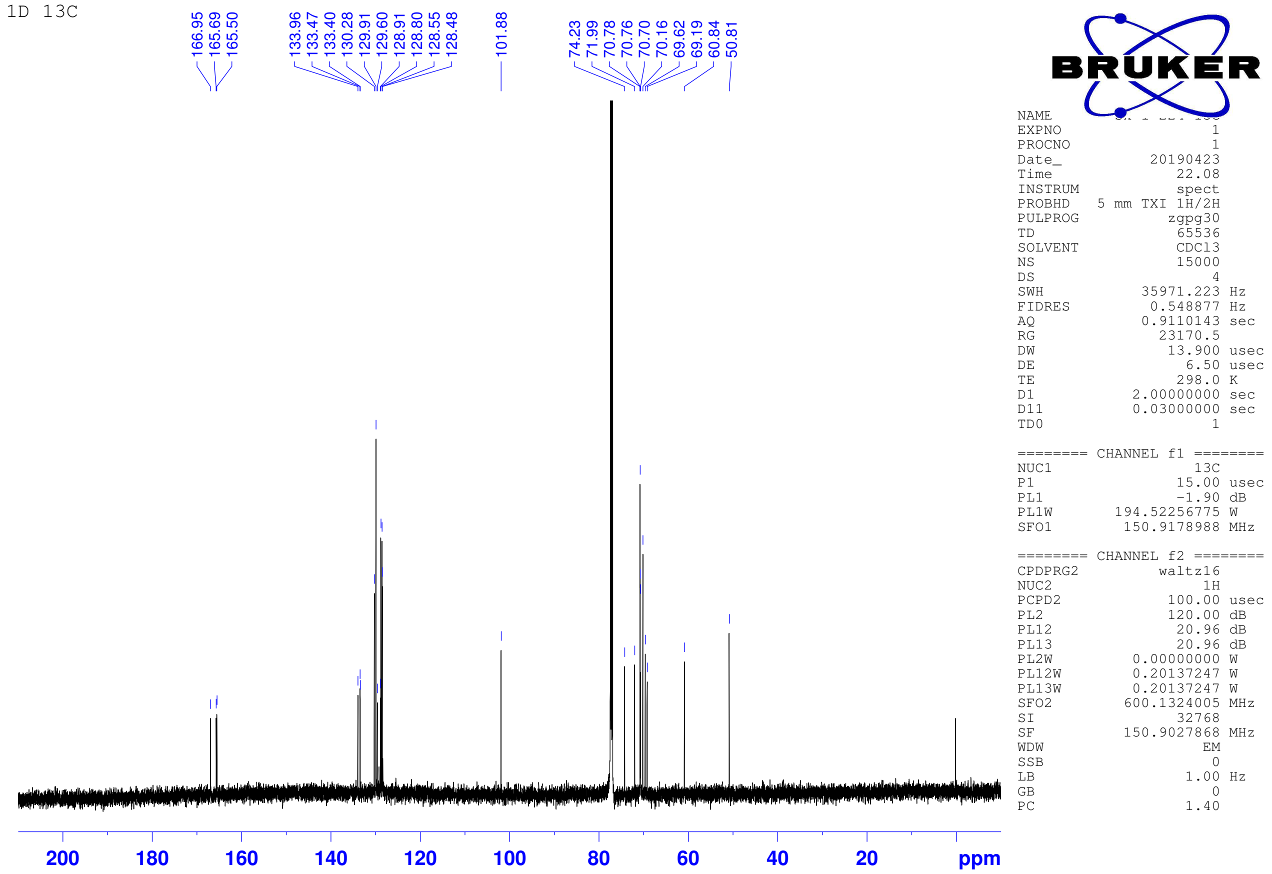

**5**: $^{1}$H NMR (600 MHz, $CDCl_3$)

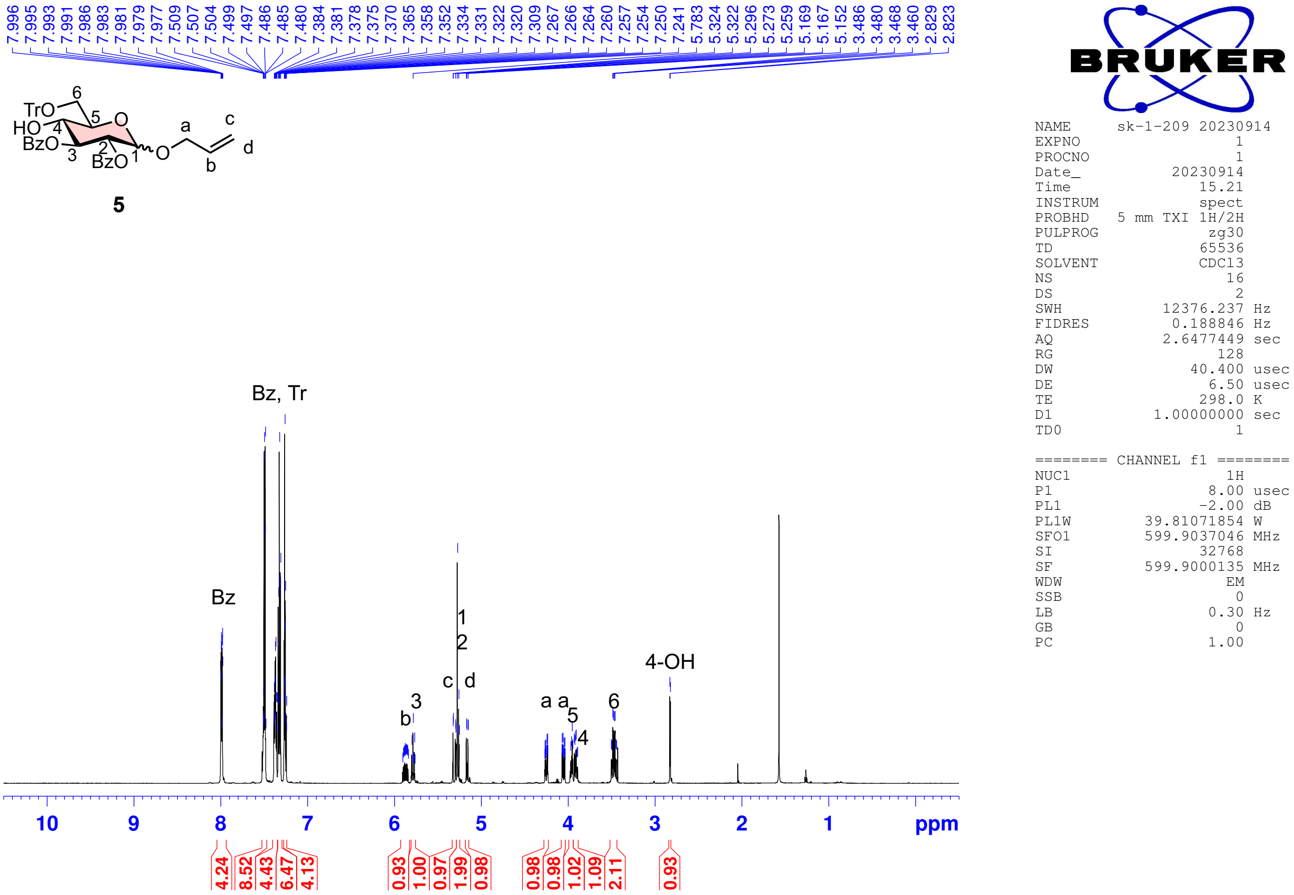


**5**: $^{13}$C NMR (150 MHz, $CDCl_3$)

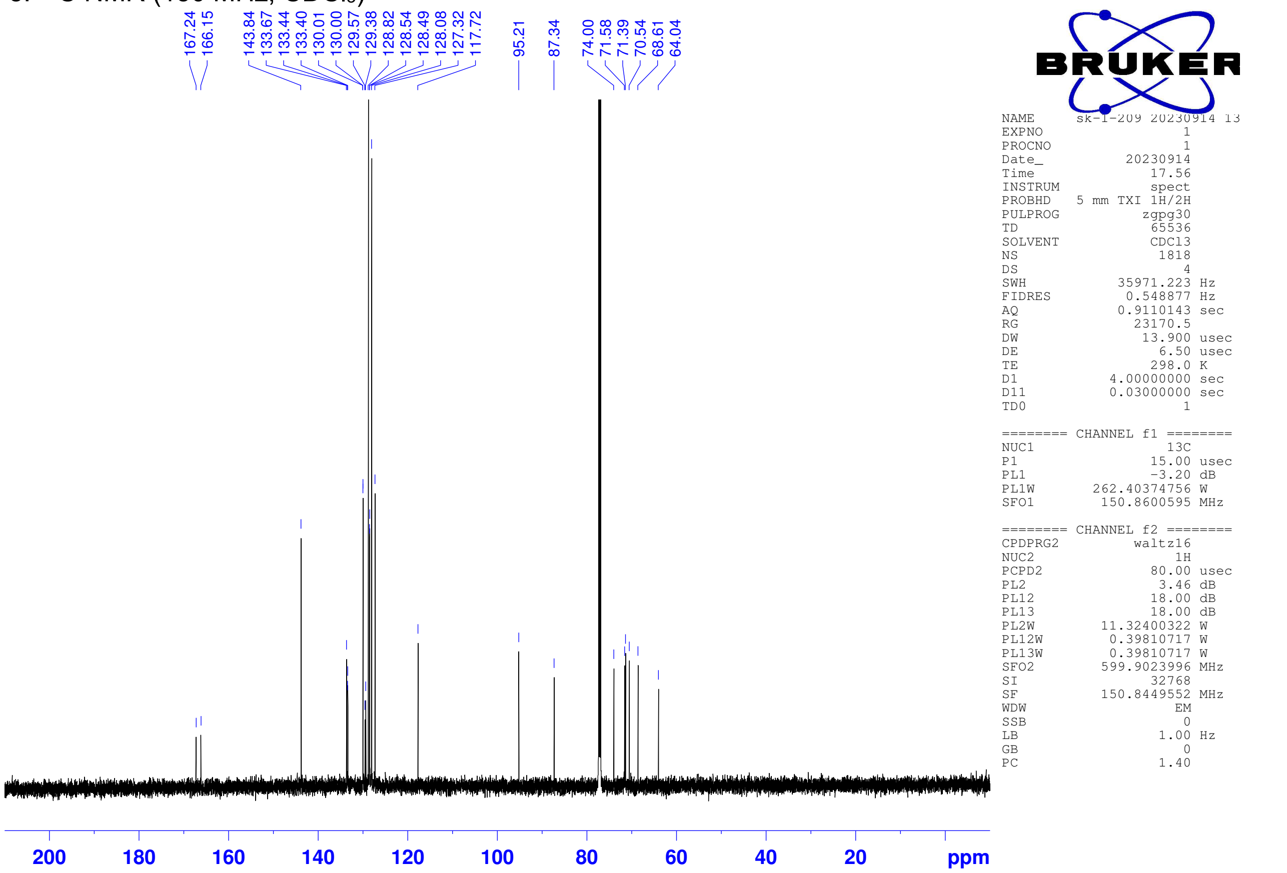

**6**: $^{1}$H NMR (600 MHz, CDCl$_3$)

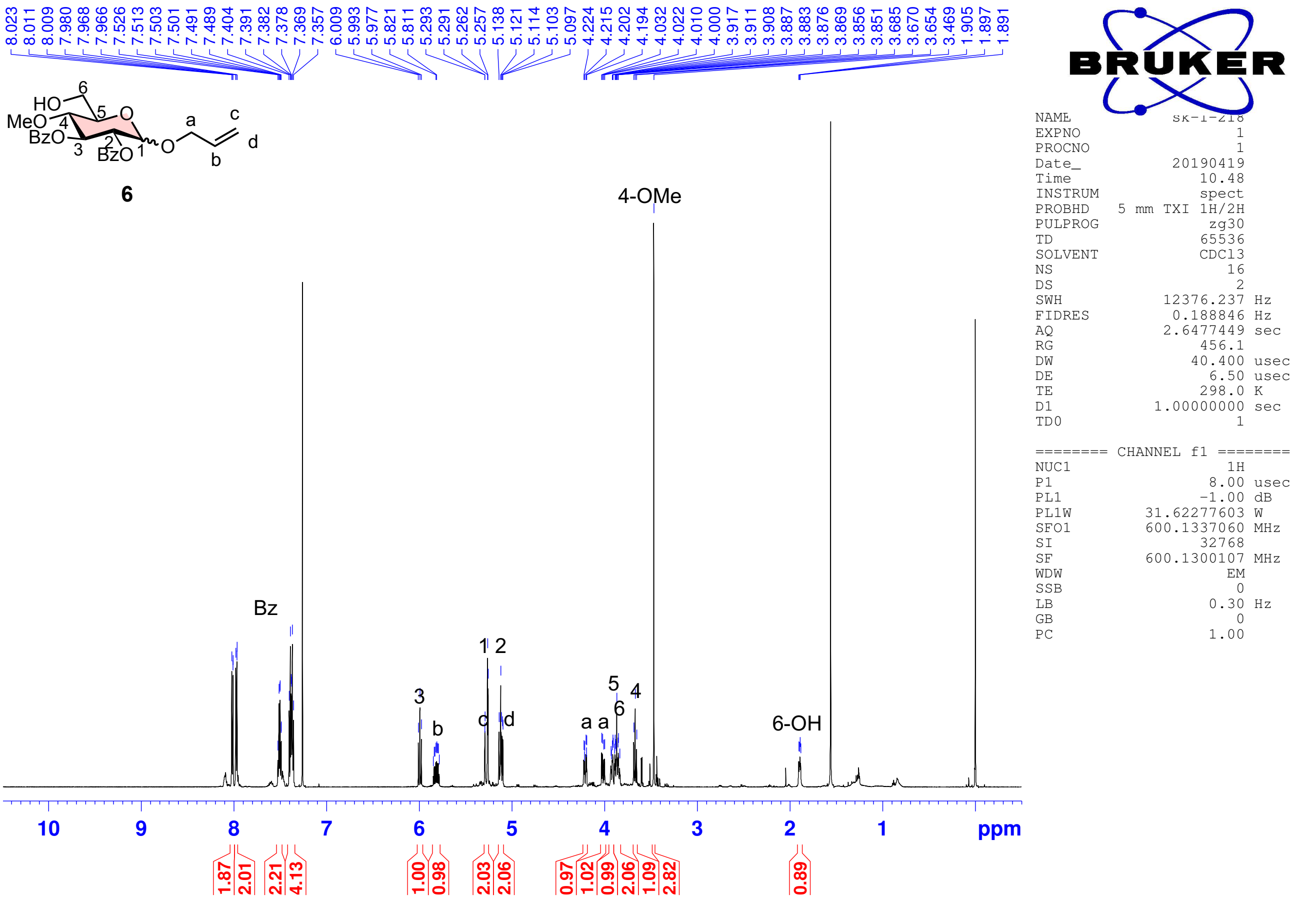


**6**: $^{13}$C NMR (150 MHz, CDCl$_3$)

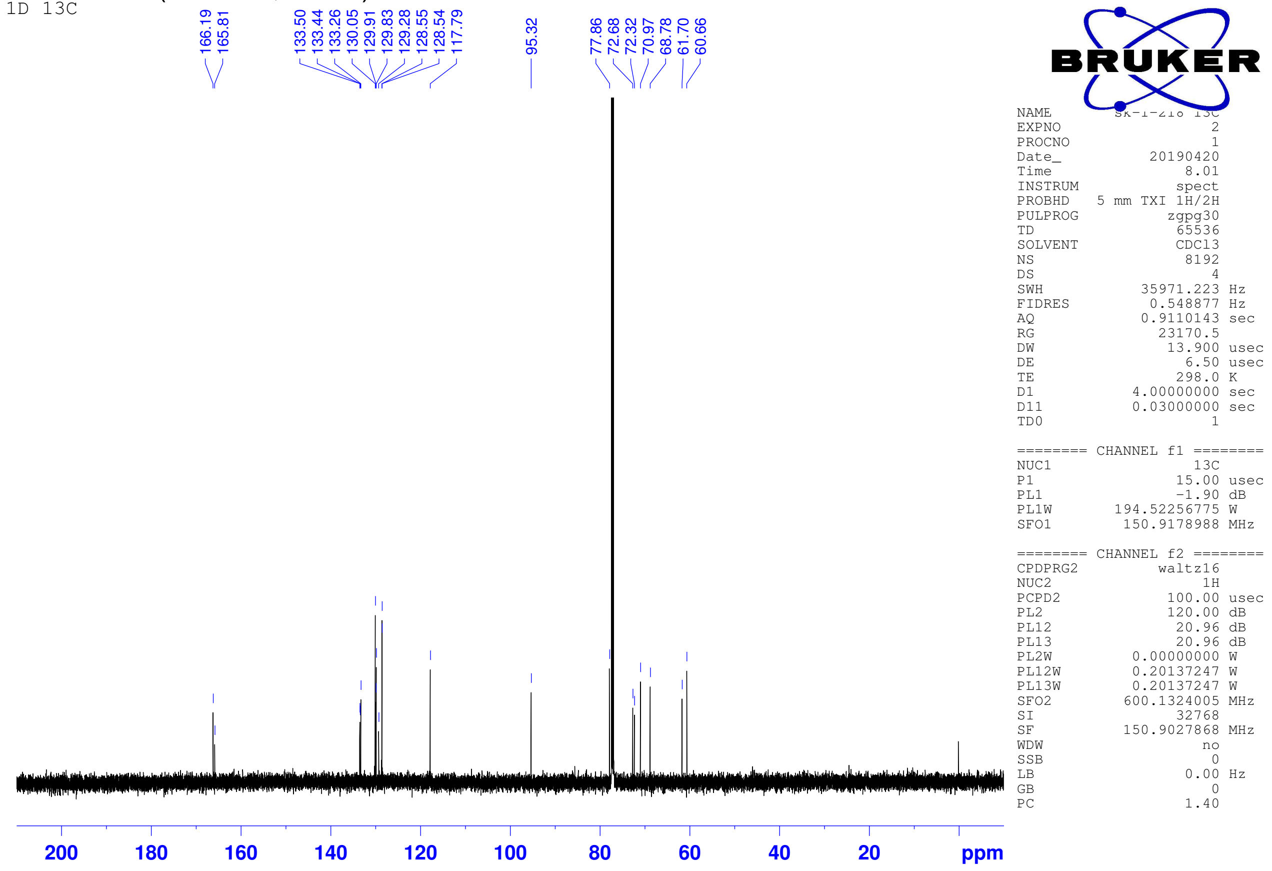

**6**: HSQC ($CDCl_3$)

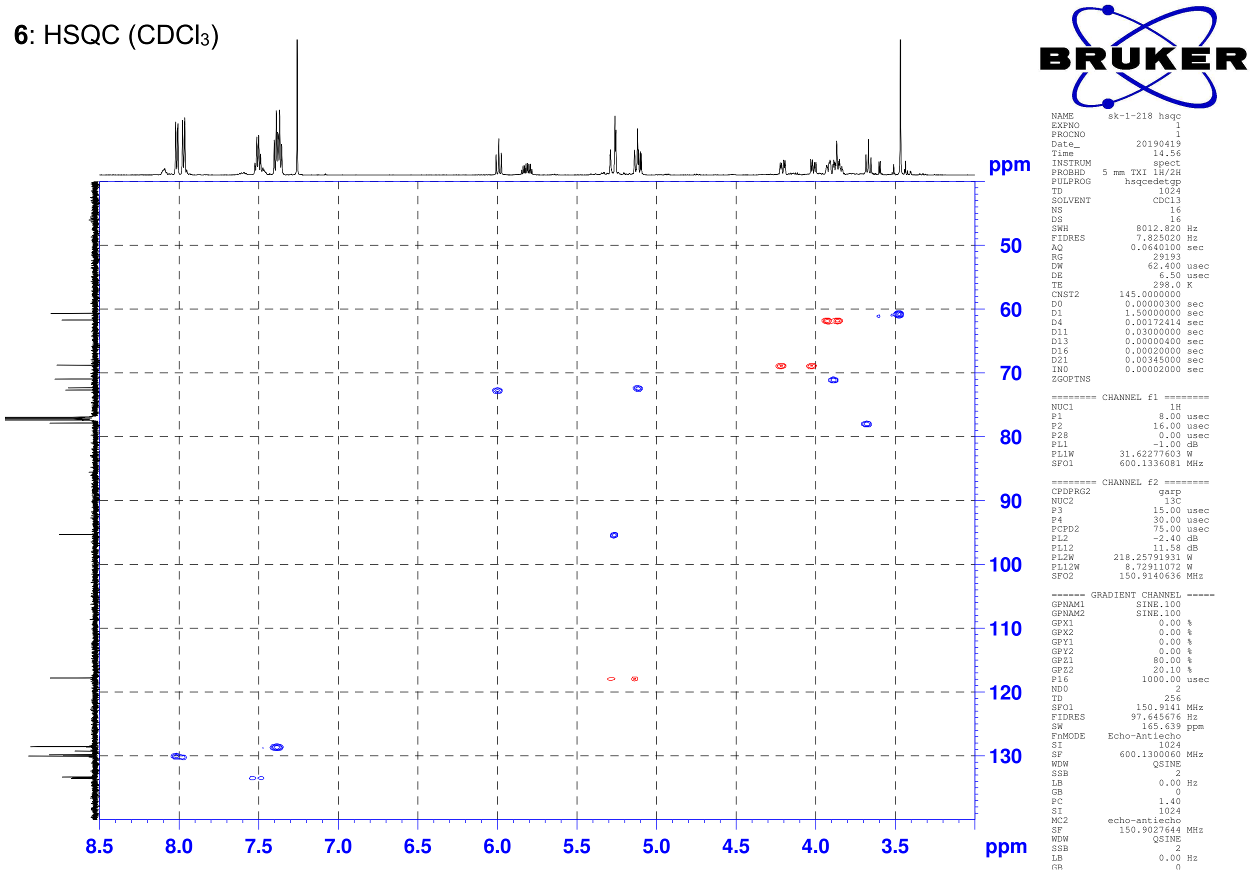


**6**: HMBC ($CDCl_3$)

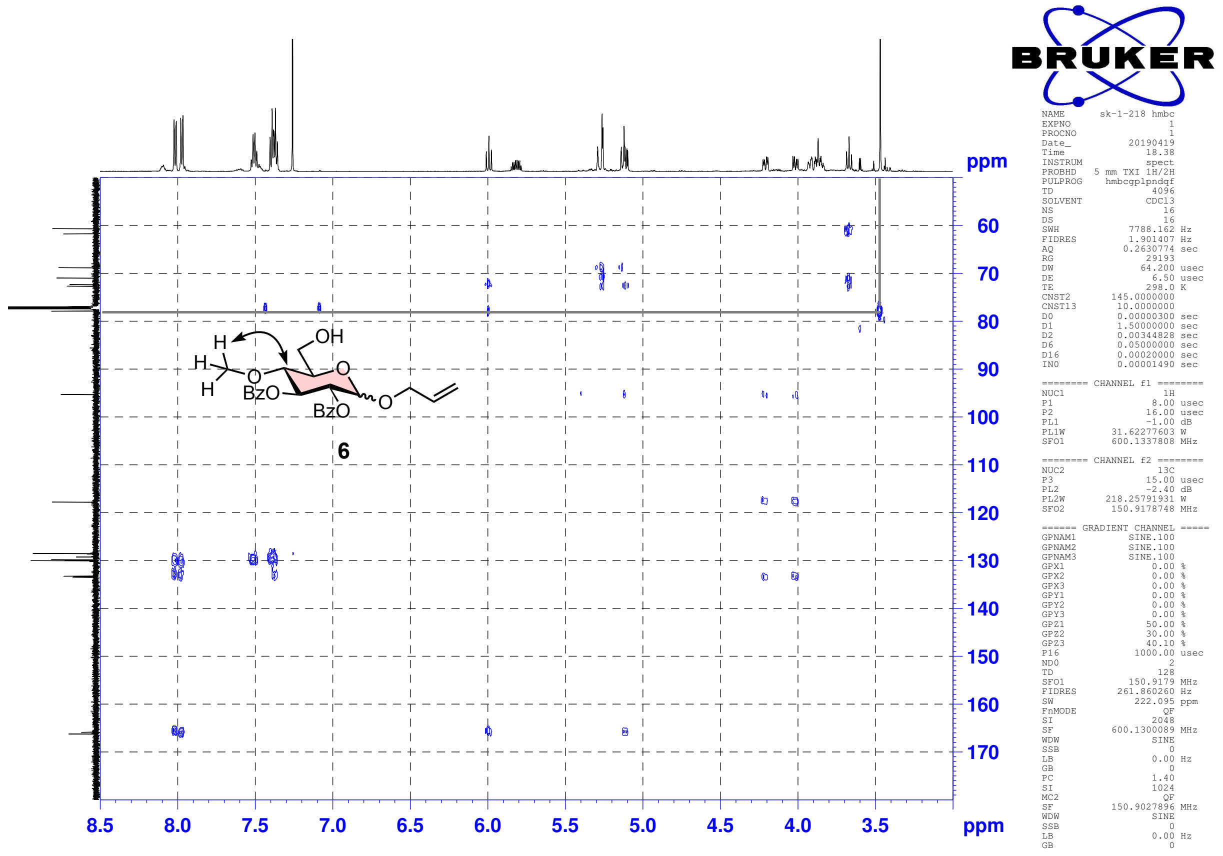

**7**: $^{1}$H NMR (600 MHz, $CDCl_3$)

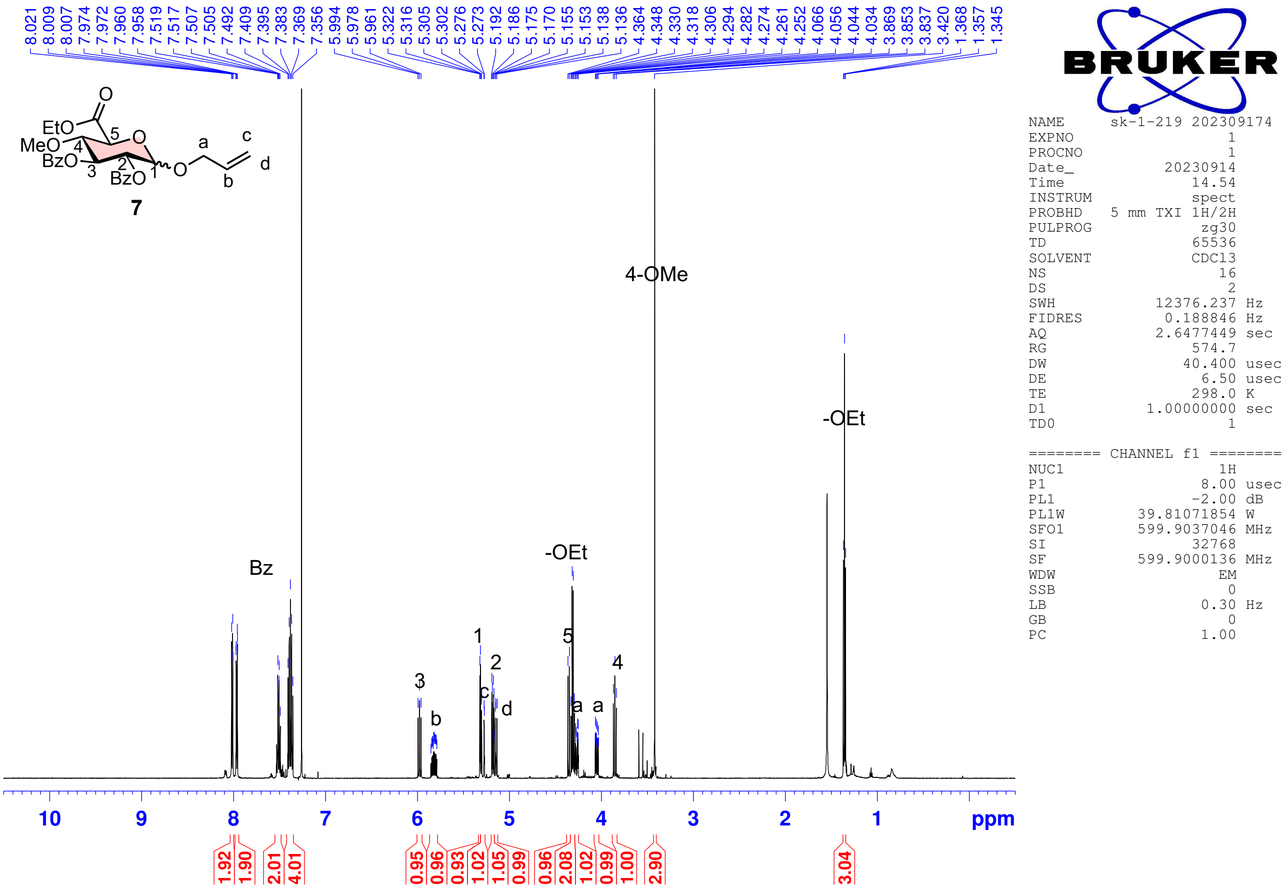


**7**: $^{13}$C NMR (150 MHz, $CDCl_3$)

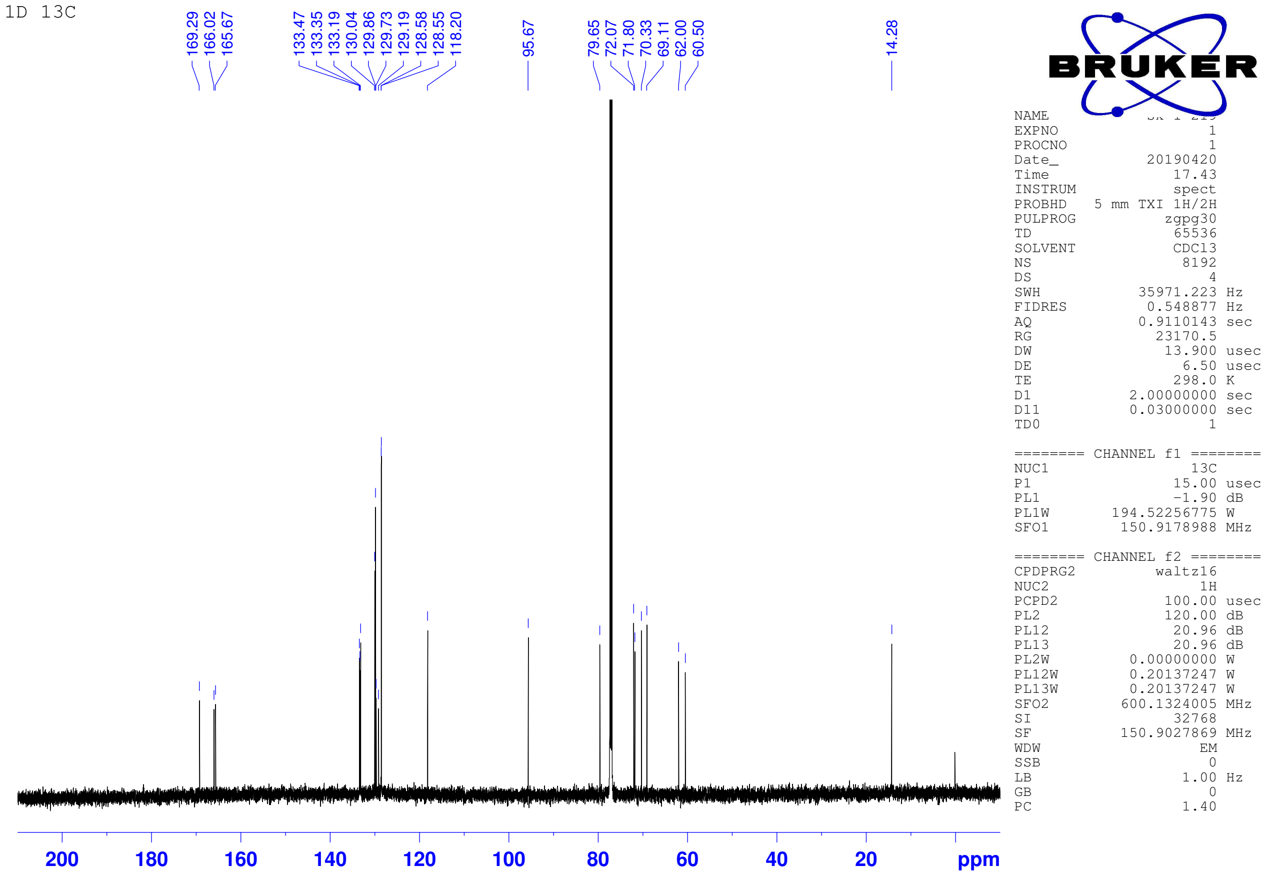

**9**: $^{1}H$ NMR (600 MHz, $CDCl_3$)

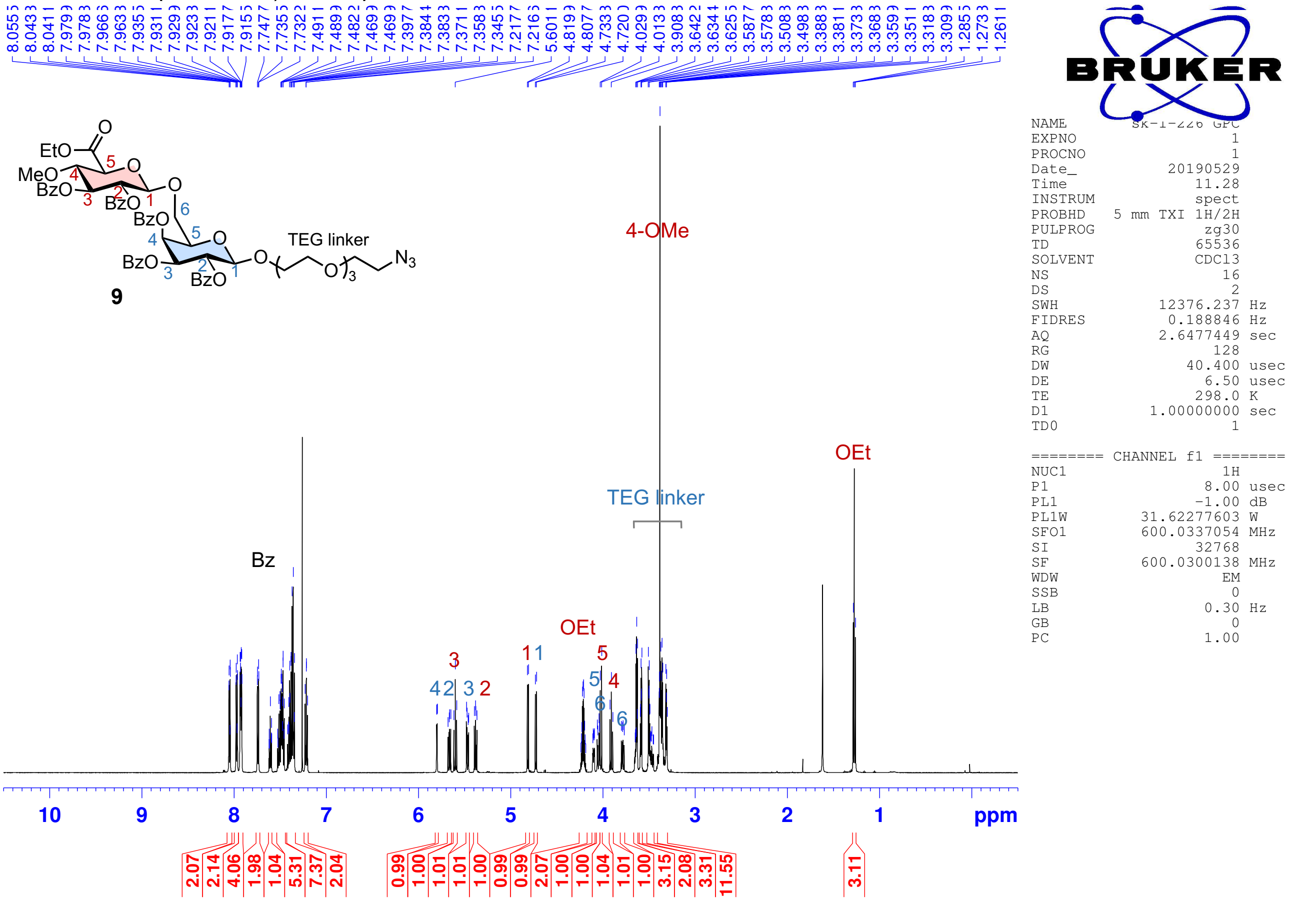


**9**: $^{13}C$ NMR (150 MHz, $CDCl_3$)

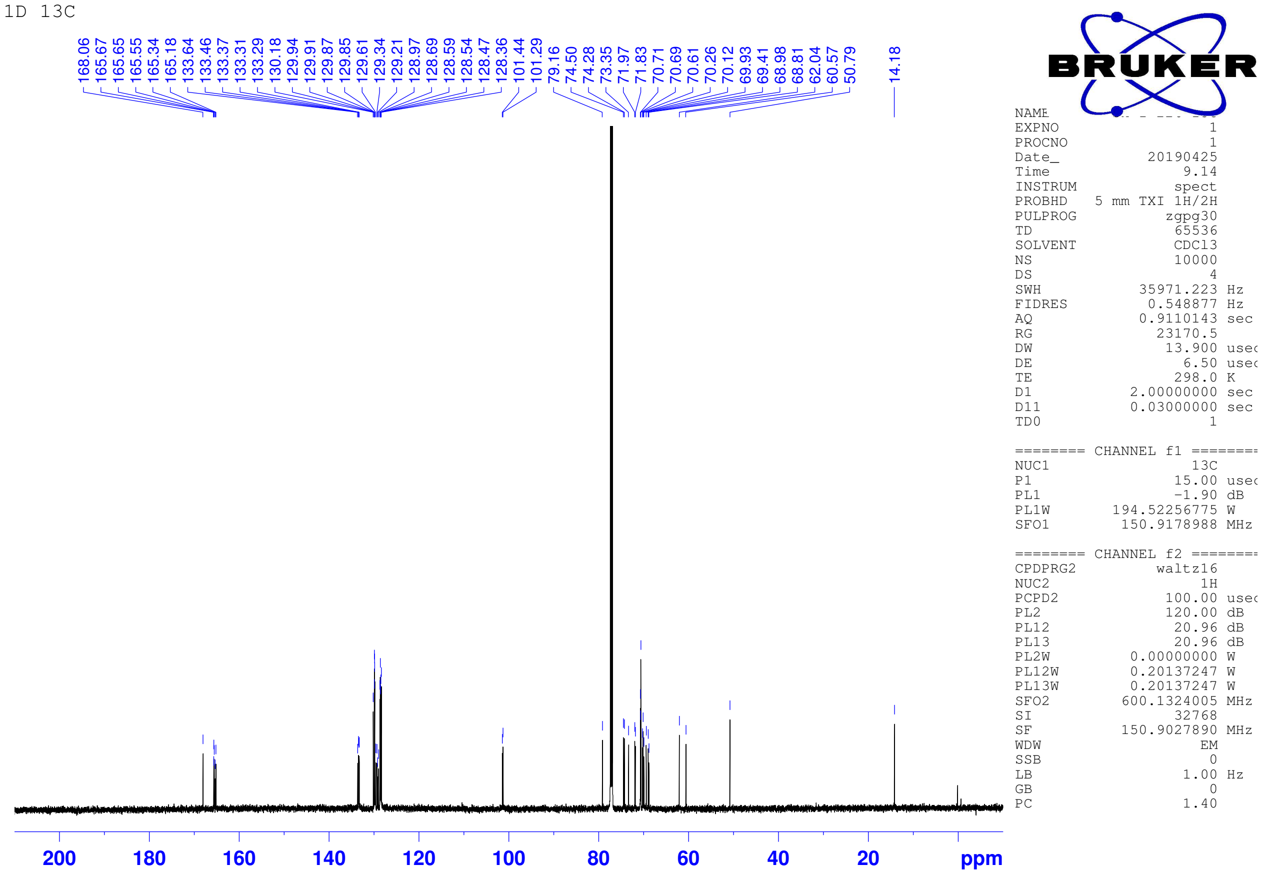

**9**: COSY ($CDCl_3$)

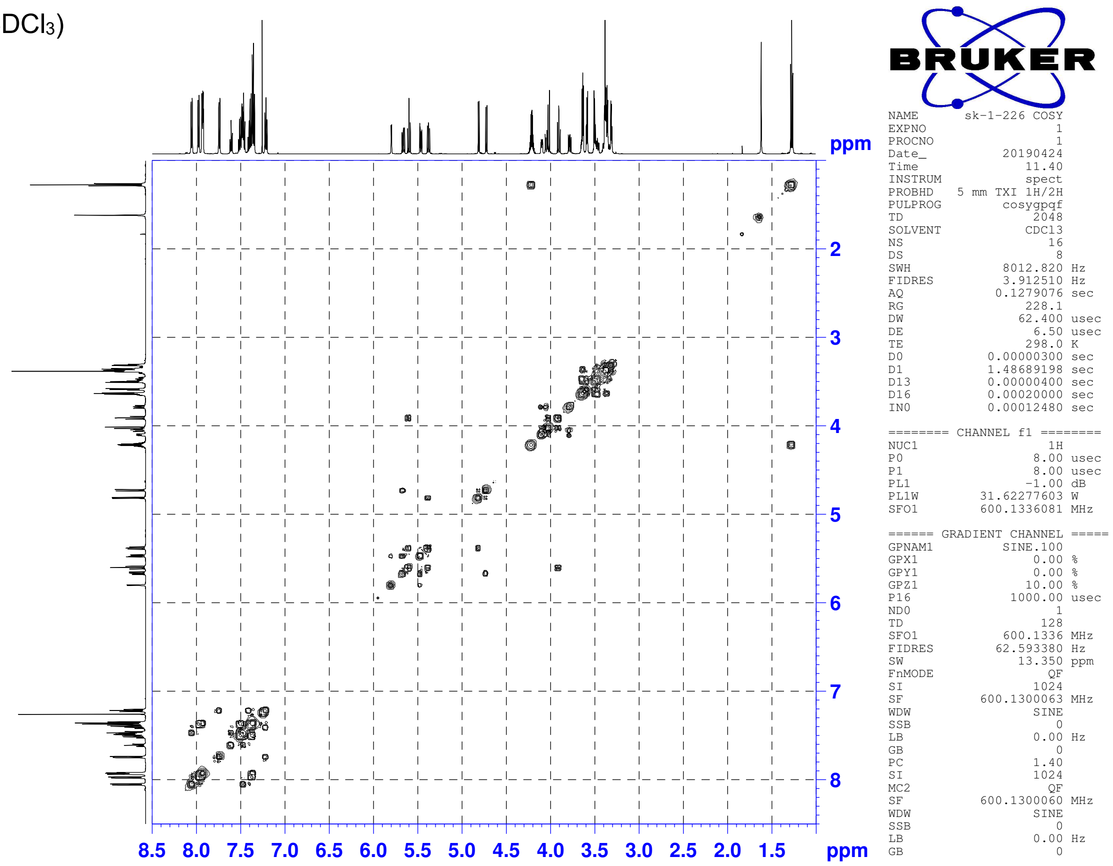


**9**: ROESY ($CDCl_3$)

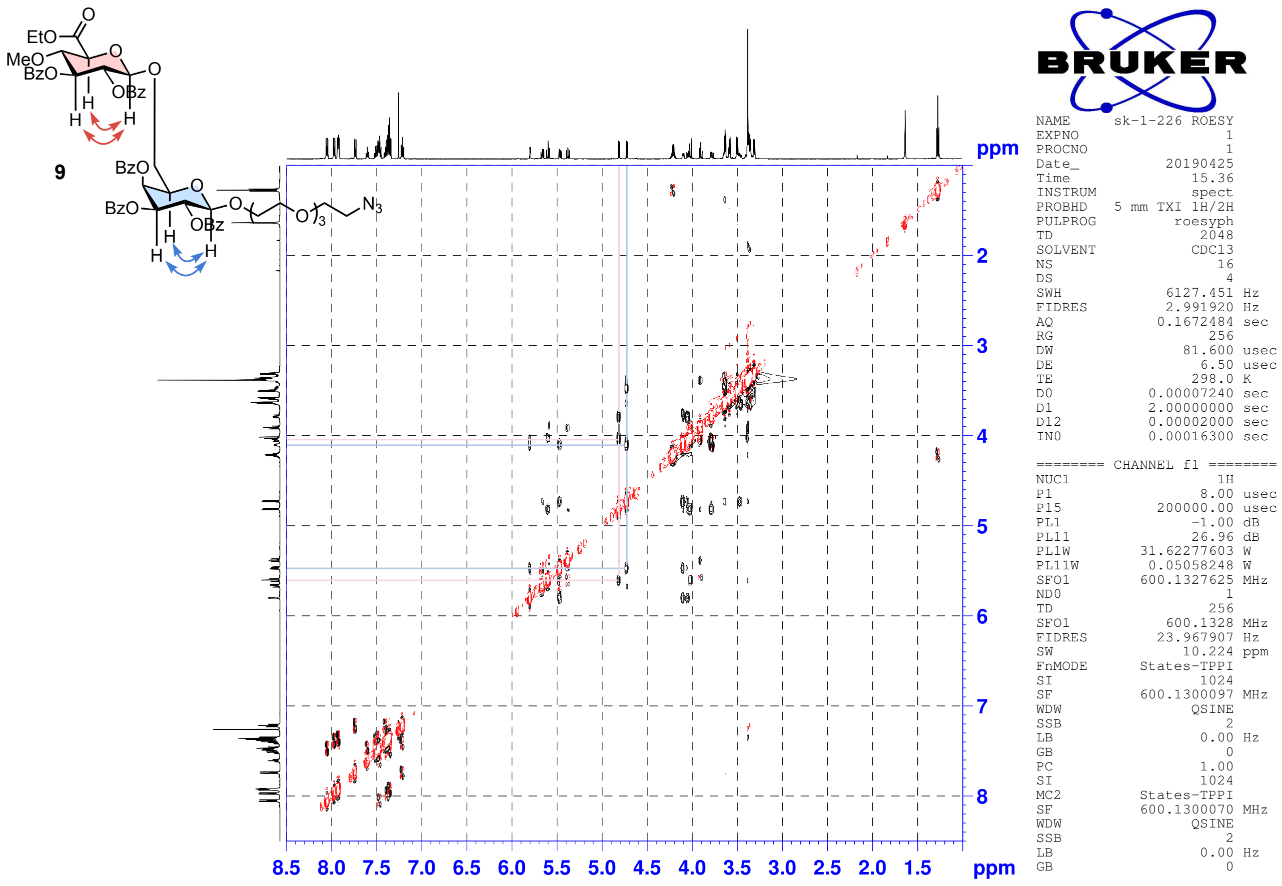

**9**: HSQC ($CDCl_3$)

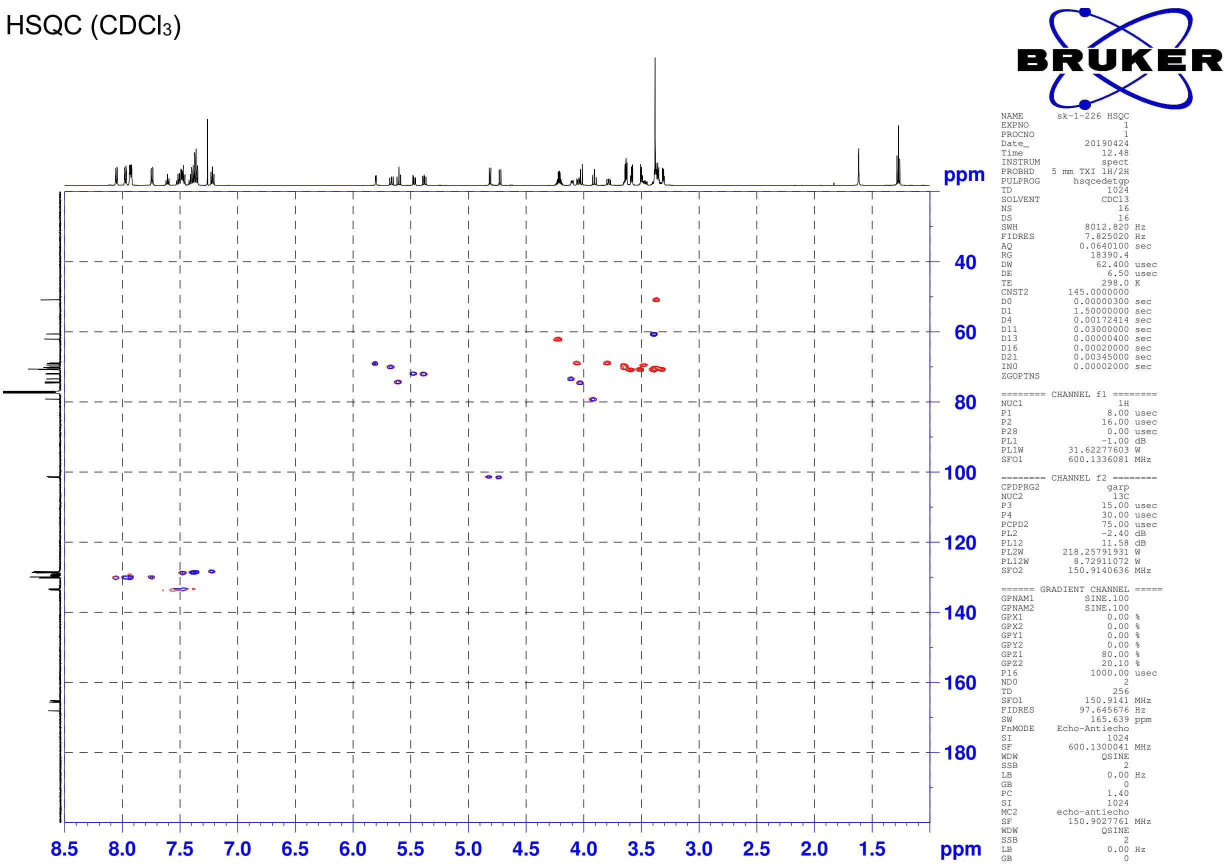


**9**: HMBC ($CDCl_3$)

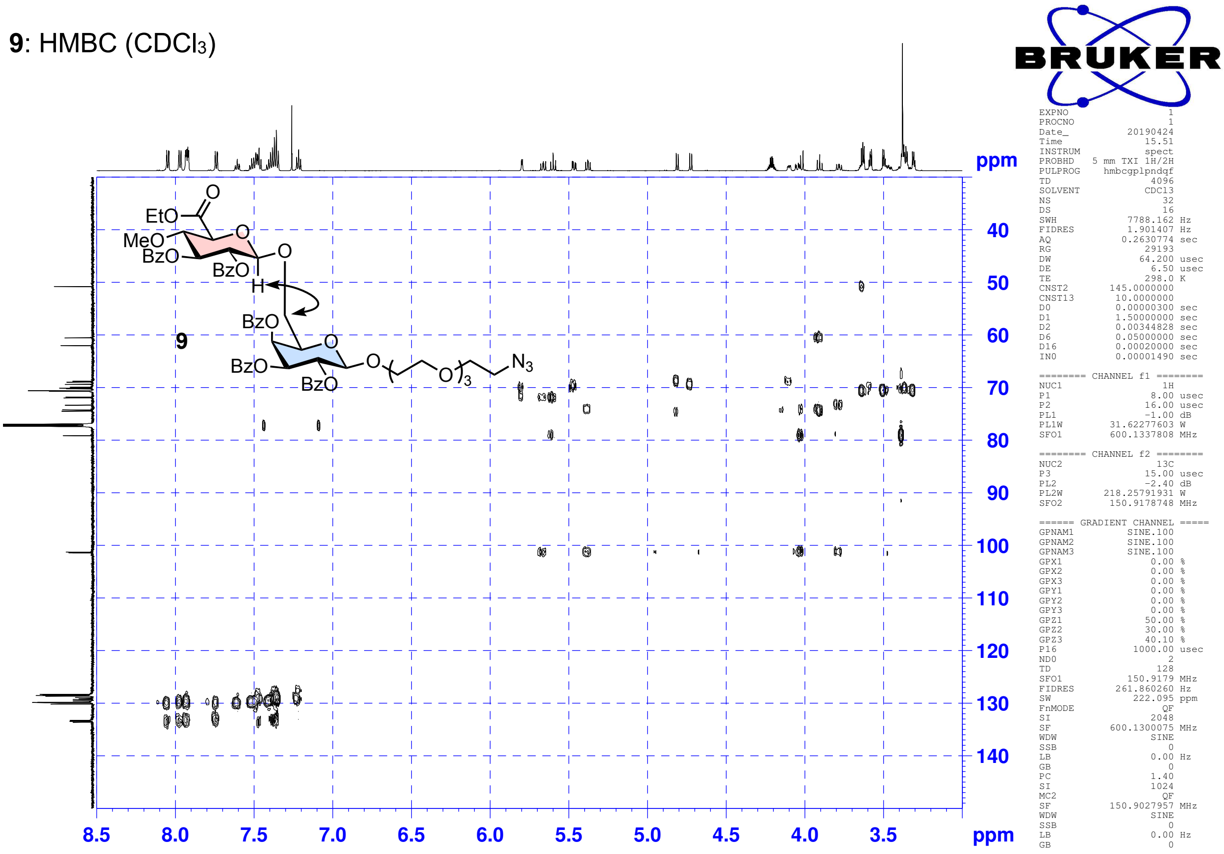

**10**: $^{1}$H NMR (600 MHz, $CDCl_3$)

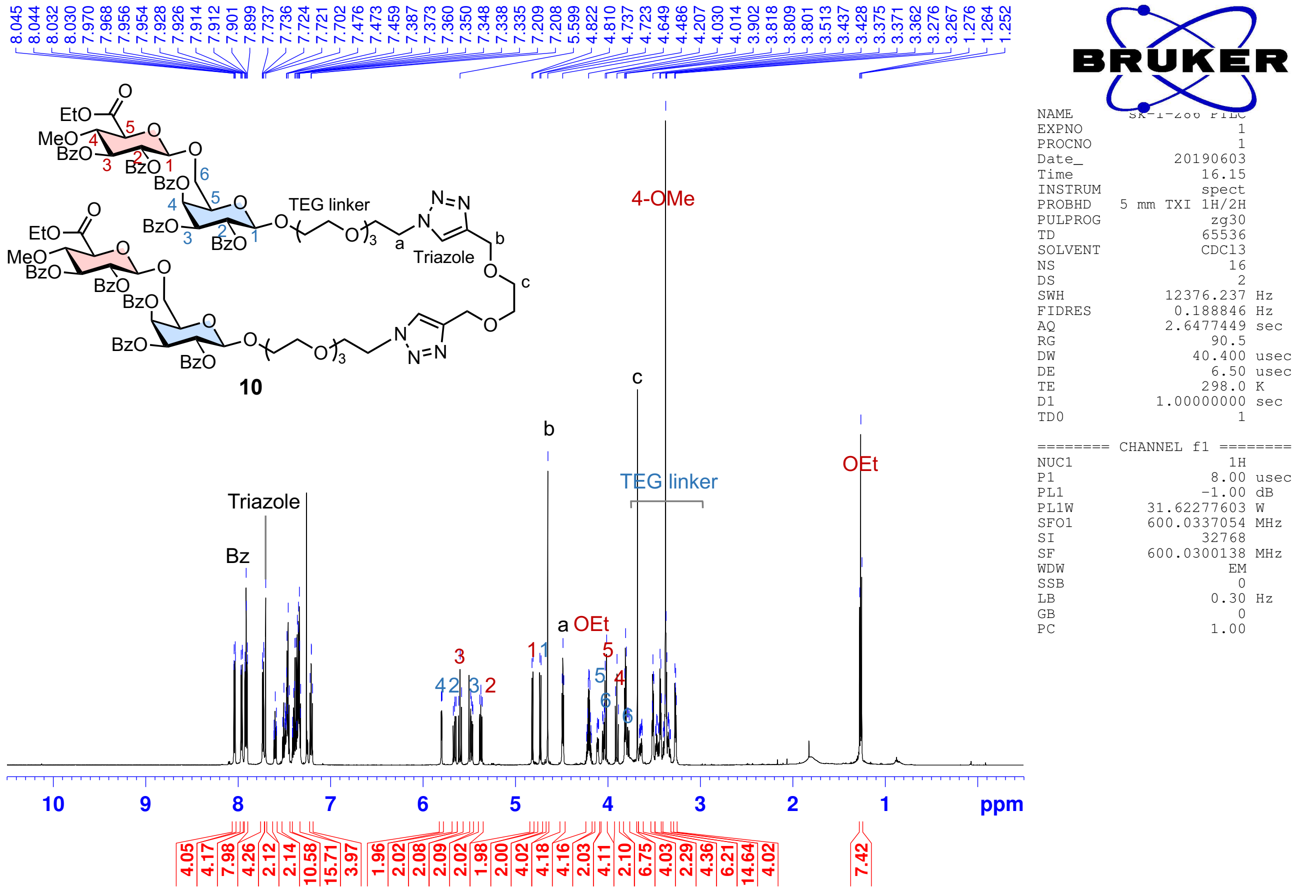


**10**: $^{13}$C NMR (150 MHz, $CDCl_3$)

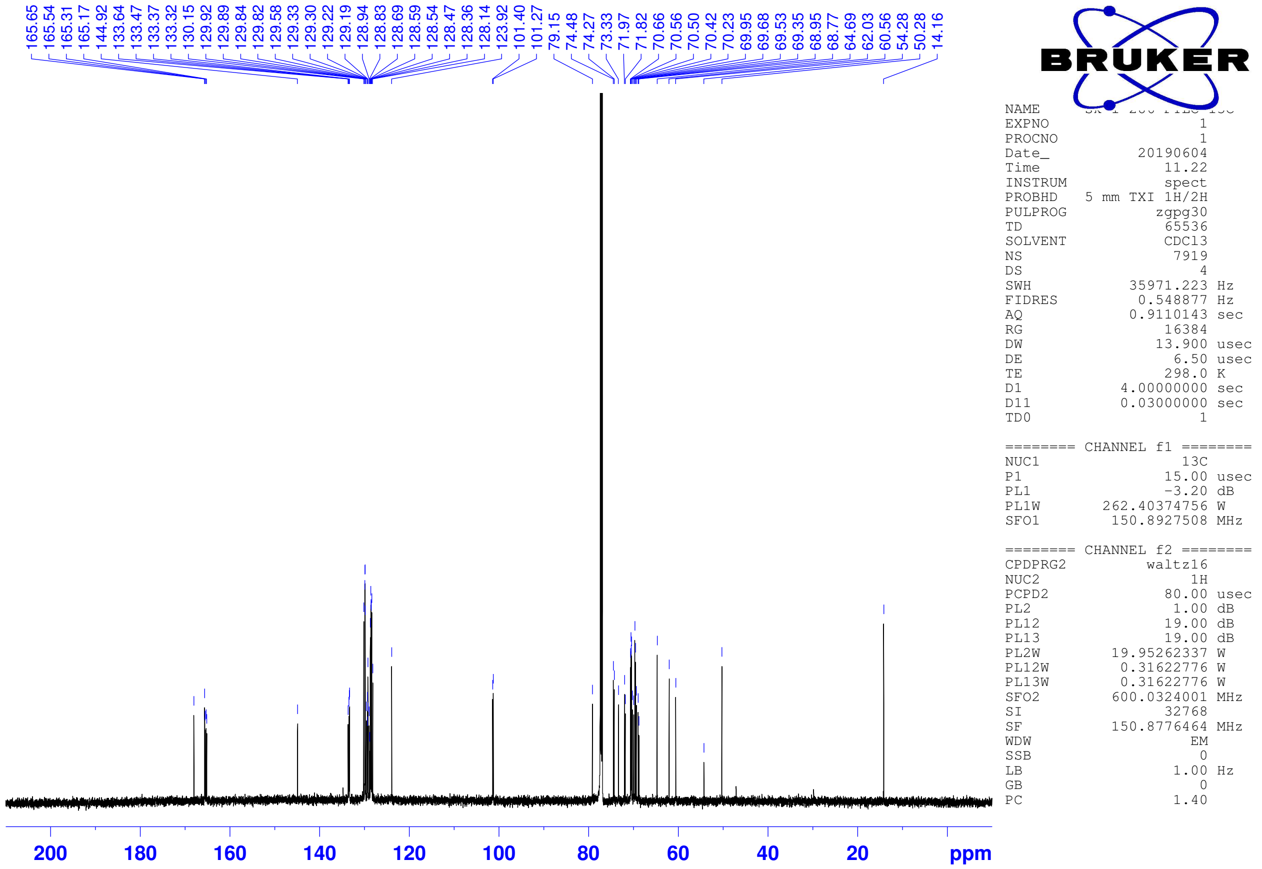

**11**: $^{1}$H NMR (600 MHz, $CDCl_3$)

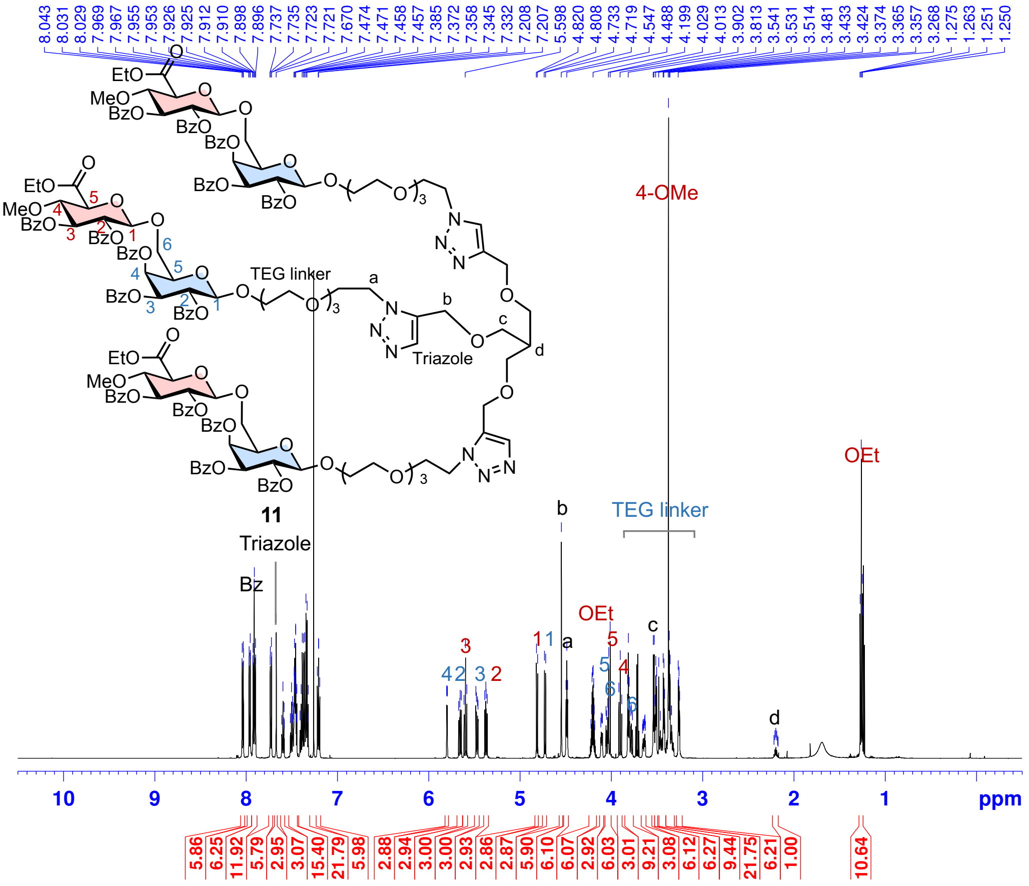


NAME sk-1-291
EXPNO 1
PROCNO 1
Date_ 20190613
Time 15.46
INSTRUM spect
PROBHD 5 mm TXI 1H/2H
PULPROG zg30
TD 65536
SOLVENT CDCl3
NS 8
DS 2
SWH 12376.237 Hz
FIDRES 0.188846 Hz
AQ 2.6477449 sec
RG 128
DW 40.400 usec
DE 6.50 usec
TE 298.0 K
D1 1.00000000 sec
TD0 1

======== CHANNEL f1 ========
NUC1 1H
P1 8.00 usec
PL1 -1.00 dB
PL1W 31.62277603 W
SFO1 600.0337054 MHz
SI 32768
SF 600.0300138 MHz
WDW EM
SSB 0
LB 0.30 Hz
GB 0
PC 1.00

**11**: $^{13}$C NMR (150 MHz, $CDCl_3$)

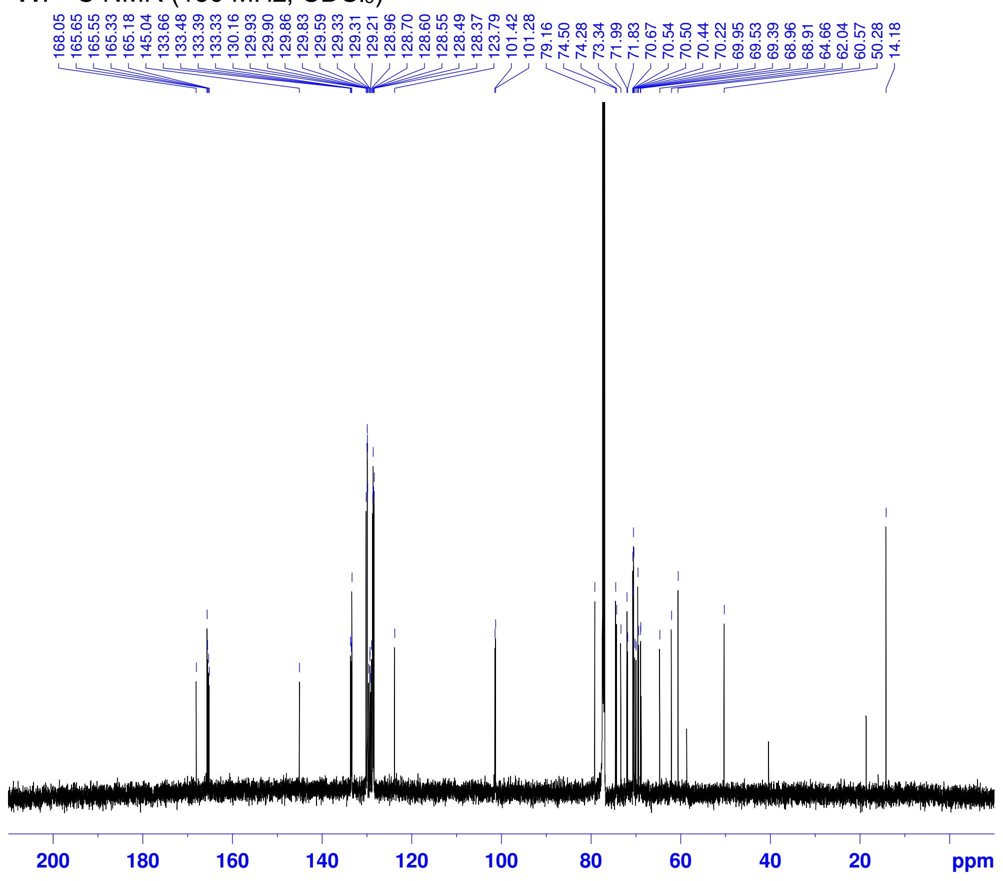


NAME sk-1-291 13C
EXPNO 1
PROCNO 1
Date_ 20190614
Time 11.30
INSTRUM spect
PROBHD 5 mm TXI 1H/2H
PULPROG zgpg30
TD 65536
SOLVENT CDCl3
NS 8192
DS 4
SWH 35971.223 Hz
FIDRES 0.548877 Hz
AQ 0.9110143 sec
RG 18390.4
DW 13.900 usec
DE 6.50 usec
TE 298.0 K
D1 4.00000000 sec
D11 0.03000000 sec
TD0 1

======== CHANNEL f1 ========
NUC1 13C
P1 15.00 usec
PL1 -3.20 dB
PL1W 262.40374756 W
SFO1 150.8927508 MHz

======== CHANNEL f2 ========
CPDPRG2 waltz16
NUC2 1H
PCPD2 80.00 usec
PL2 1.00 dB
PL12 19.00 dB
PL13 19.00 dB
PL2W 19.95262337 W
PL12W 0.31622776 W
PL13W 0.31622776 W
SFO2 600.0324001 MHz
SI 32768
SF 150.8776442 MHz
WDW EM
SSB 0
LB 1.00 Hz
GB 0
PC 1.40

**12** $^1$H NMR (600 MHz, $CDCl_3$)

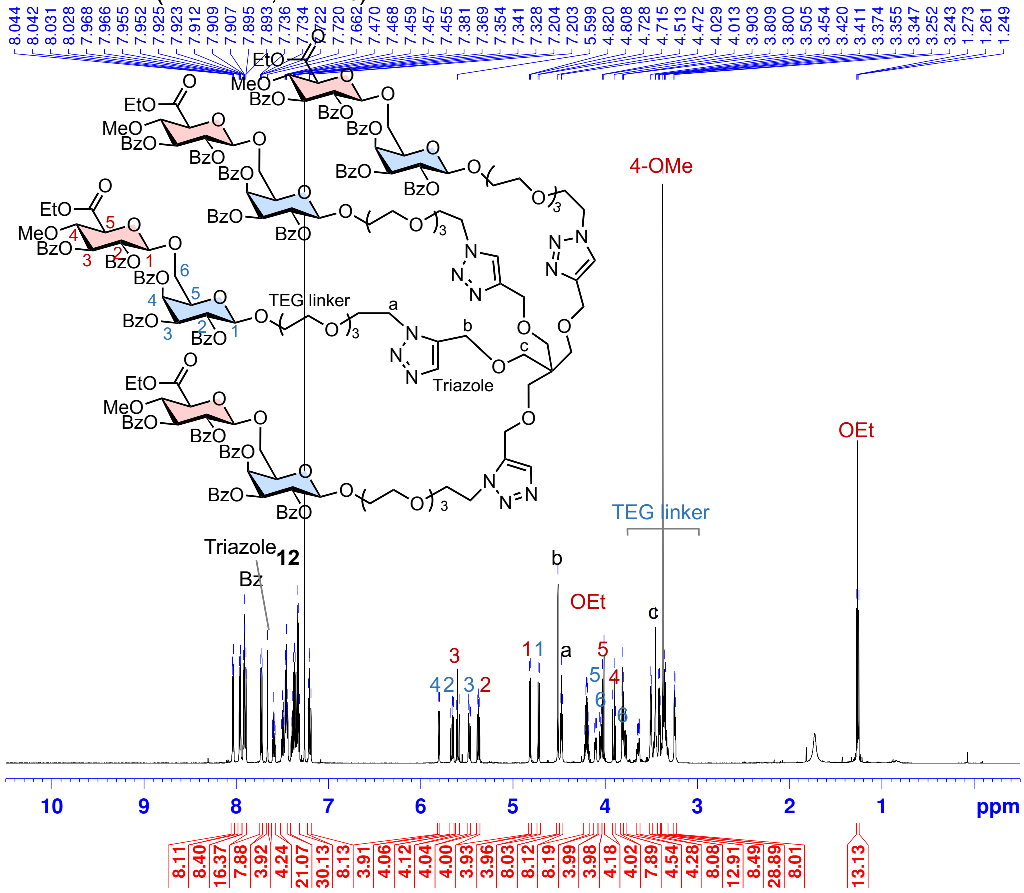


| | |
|---|---|
| NAME | sk-1-295 |
| EXPNO | 1 |
| PROCNO | 1 |
| Date_ | 20190614 |
| Time | 19.03 |
| INSTRUM | spect |
| PROBHD | 5 mm TXI 1H/2H |
| PULPROG | zg30 |
| TD | 65536 |
| SOLVENT | CDCl3 |
| NS | 16 |
| DS | 2 |
| SWH | 12376.237 Hz |
| FIDRES | 0.188846 Hz |
| AQ | 2.6477449 sec |
| RG | 128 |
| DW | 40.400 usec |
| DE | 6.50 usec |
| TE | 298.0 K |
| D1 | 1.00000000 sec |
| TD0 | 1 |
| ======== CHANNEL f1 ======== | |
| NUC1 | 1H |
| P1 | 8.00 usec |
| PL1 | -1.00 dB |
| PL1W | 31.62277603 W |
| SFO1 | 600.0337054 MHz |
| SI | 32768 |
| SF | 600.0300138 MHz |
| WDW | EM |
| SSB | 0 |
| LB | 0.30 Hz |
| GB | 0 |
| PC | 1.00 |

**12**: $^{13}$C NMR (150 MHz, $CDCl_3$)

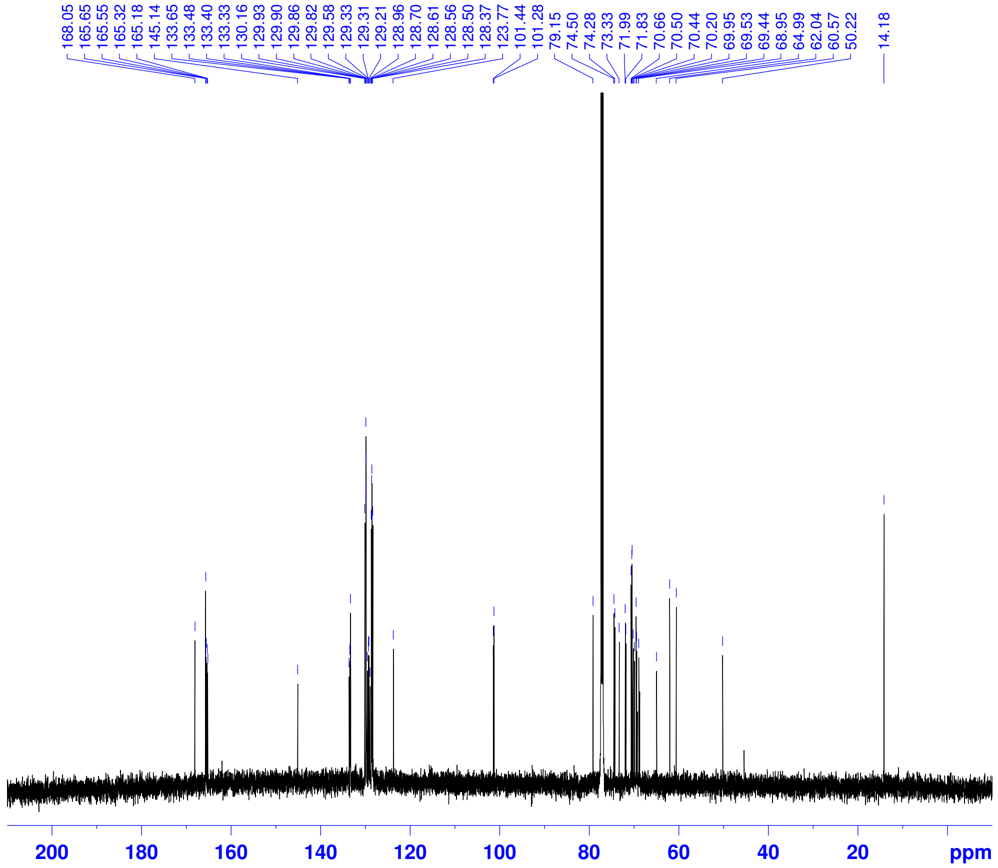

| | |
|---|---|
| NAME | sk-1-295 13C |
| EXPNO | 1 |
| PROCNO | 1 |
| Date_ | 20190615 |
| Time | 13.12 |
| INSTRUM | spect |
| PROBHD | 5 mm TXI 1H/2H |
| PULPROG | zgpg30 |
| TD | 65536 |
| SOLVENT | CDCl3 |
| NS | 8192 |
| DS | 4 |
| SWH | 35971.223 Hz |
| FIDRES | 0.548877 Hz |
| AQ | 0.9110143 sec |
| RG | 20642.5 |
| DW | 13.900 usec |
| DE | 6.50 usec |
| TE | 298.0 K |
| D1 | 4.00000000 sec |
| D11 | 0.03000000 sec |
| TD0 | 1 |
| ======== CHANNEL f1 ======== | |
| NUC1 | 13C |
| P1 | 15.00 usec |
| PL1 | -3.20 dB |
| PL1W | 262.40374756 W |
| SFO1 | 150.8927508 MHz |
| ======== CHANNEL f2 ======== | |
| CPDPRG2 | waltz16 |
| NUC2 | 1H |
| PCPD2 | 80.00 usec |
| PL2 | 1.00 dB |
| PL12 | 19.00 dB |
| PL13 | 19.00 dB |
| PL2W | 19.95262337 W |
| PL12W | 0.31622776 W |
| PL13W | 0.31622776 W |
| SFO2 | 600.0324001 MHz |
| SI | 32768 |
| SF | 150.8776440 MHz |
| WDW | EM |
| SSB | 0 |
| LB | 1.00 Hz |
| GB | 0 |
| PC | 1.40 |

**AMOR dimer**: $^{1}$H NMR (600 MHz, $D_2O$)

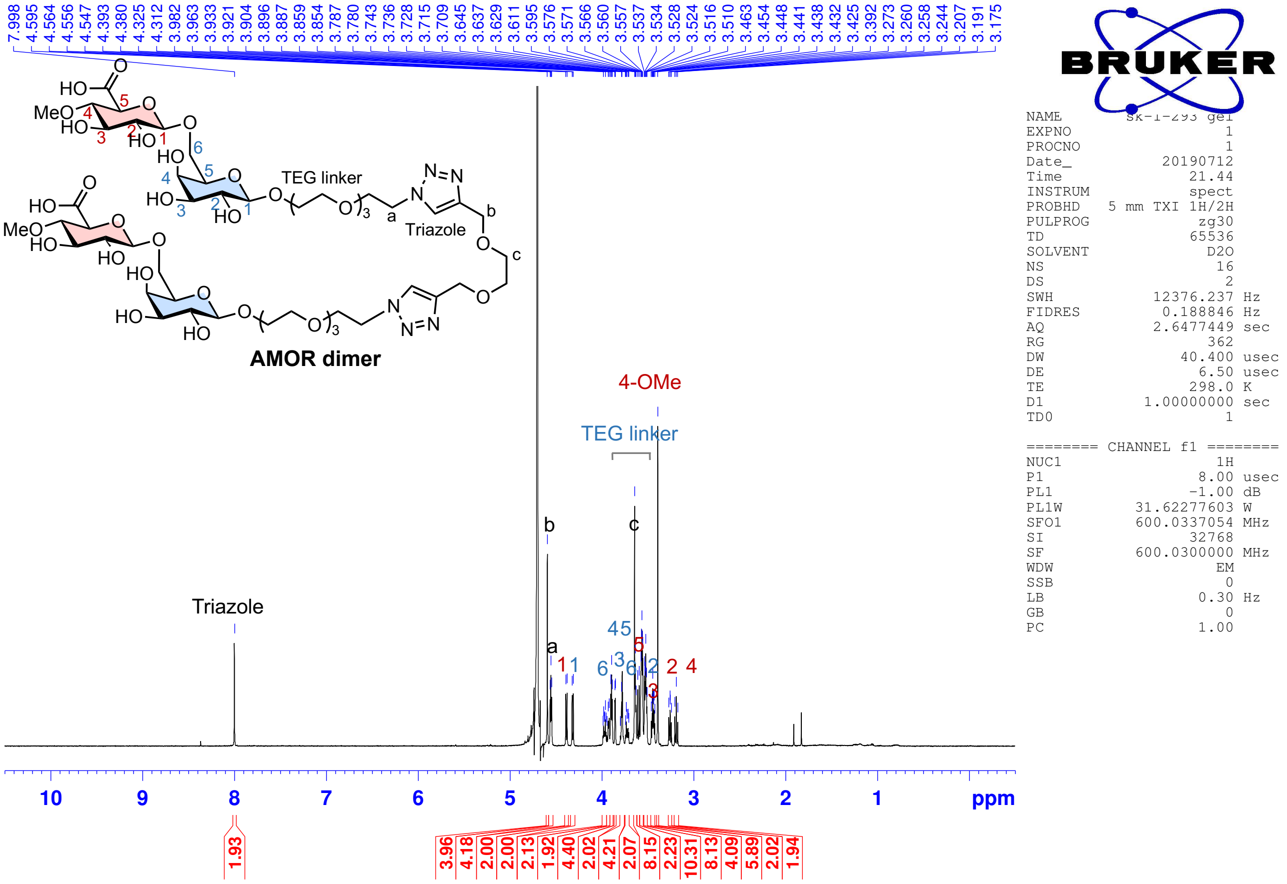


**AMOR trimer**: $^{1}$H NMR (600 MHz, $D_2O$)

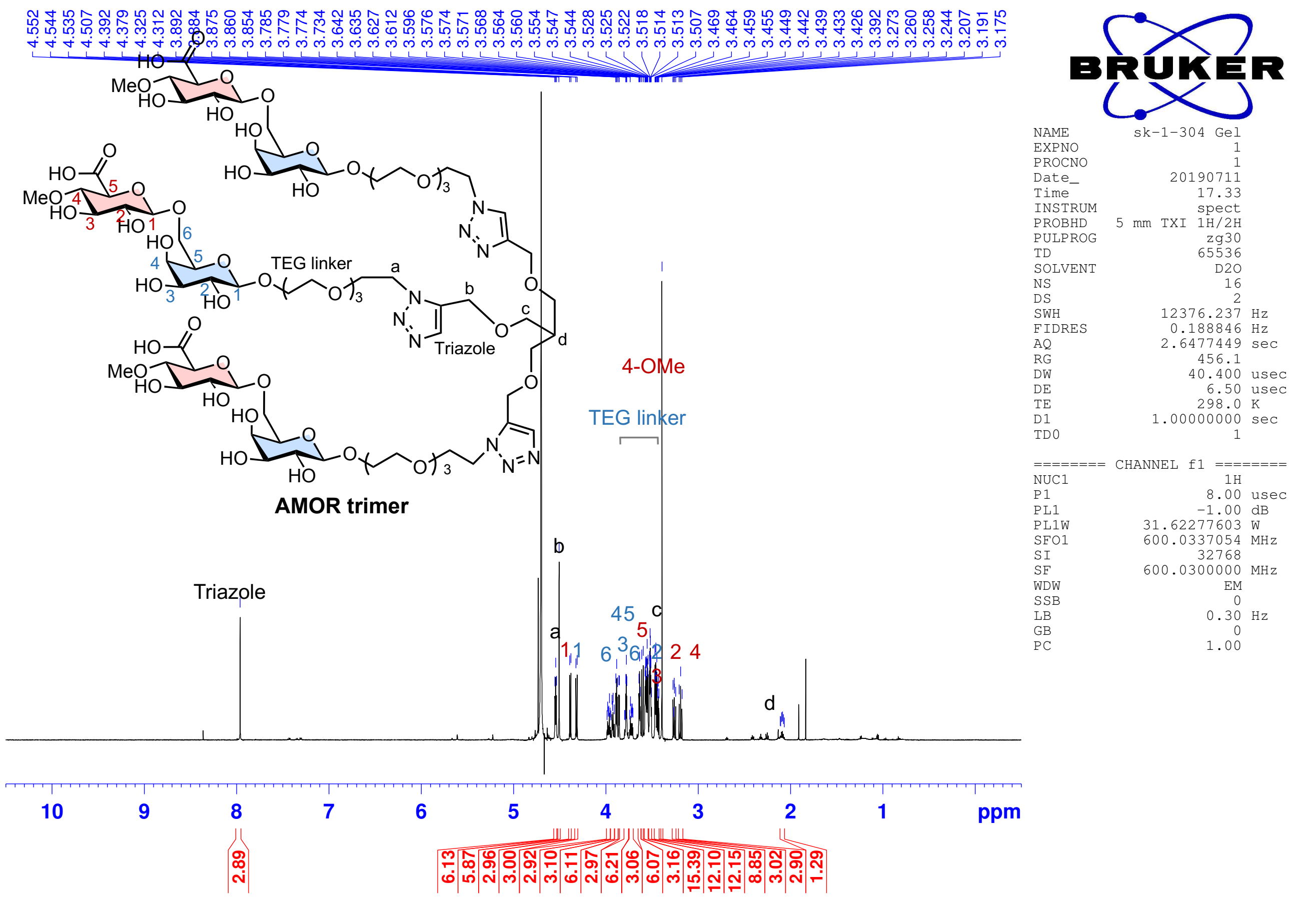

**AMOR tetramer**: $^{1}H$ NMR (600 MHz, $D_2O$)

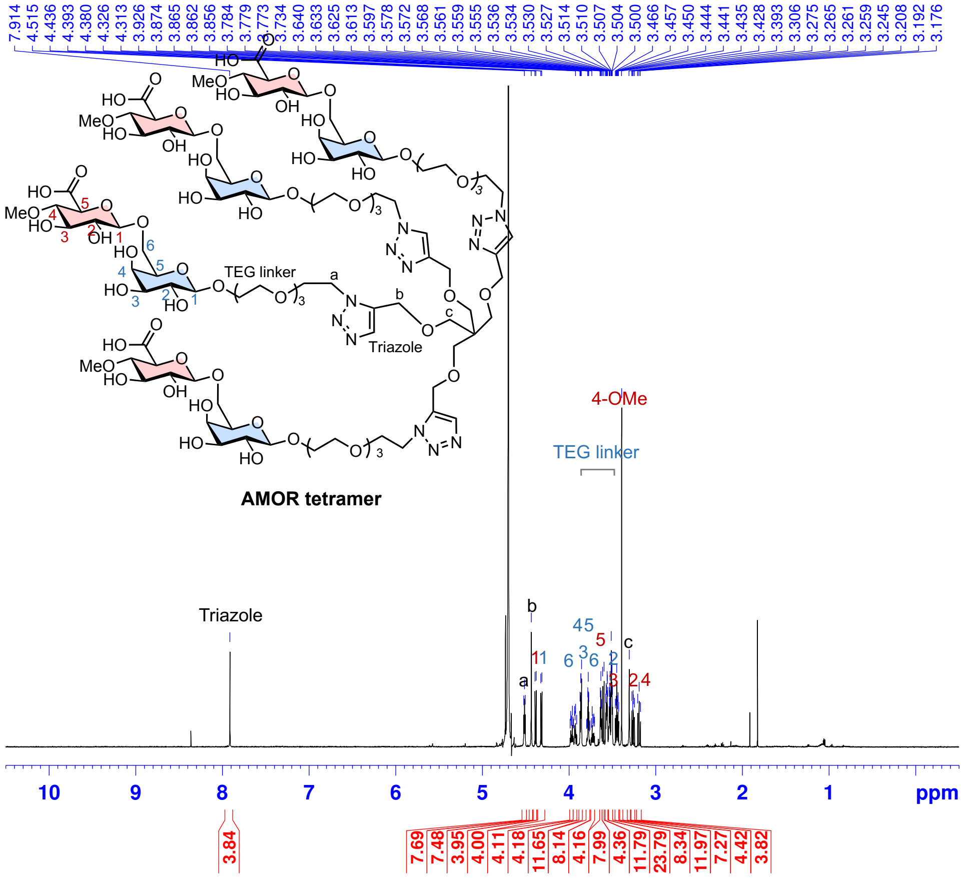


| | |
|---|---|
| NAME | sk-1-305 gel |
| EXPNO | 1 |
| PROCNO | 1 |
| Date_ | 20190712 |
| Time | 21.35 |
| INSTRUM | spect |
| PROBHD | 5 mm TXI 1H/2H |
| PULPROG | zg30 |
| TD | 65536 |
| SOLVENT | D2O |
| NS | 16 |
| DS | 2 |
| SWH | 12376.237 Hz |
| FIDRES | 0.188846 Hz |
| AQ | 2.6477449 sec |
| RG | 512 |
| DW | 40.400 usec |
| DE | 6.50 usec |
| TE | 298.0 K |
| D1 | 1.00000000 sec |
| TD0 | 1 |
| ======== CHANNEL f1 ======== | |
| NUC1 | 1H |
| P1 | 8.00 usec |
| PL1 | -1.00 dB |
| PL1W | 31.62277603 W |
| SFO1 | 600.0337054 MHz |
| SI | 32768 |
| SF | 600.0300000 MHz |
| WDW | EM |
| SSB | 0 |
| LB | 0.30 Hz |
| GB | 0 |
| PC | 1.00 |

#### 4. Bioassay

**Plant Materials and Growth conditions**

Torenia fournieri cv 'blue and white' (Sakata no Tane) were grown on soil at 28 °C with a 16-h photoperiod [approximately 150 μmol / ($m^{-2}s^{-1}$)].

**AMOR activity of synthesized AMOR and AMOR oligomers**

AMOR activity was measured using AMOR assay, which have been previously reported[4,5] (Mizukami et al, 2016; Jiao et al, 2017). The center of an agar plate of modified Nitsch's medium in a glass-bottom dish (D210402; Matsunami) was cut into an ~18 x 22 mm2 square and the agar block was removed. In this space, growth medium [modified Nitsch's medium containing 13% polyethylene glycol 4000 (w/v), and 1% sucrose (w/v)] containing AMOR (TCI) or multivalent AMORs with 1.5% ultla-low gelling agarose (agarose type IX-A; Sigma) were spread on the glass botton to form a thin layer of agar and the cut end of 15-mm-long hand-pollinated style was embedded before solidification. This agar plate was incubated in the dark at 28 °C for 16 h. After pollen tube incubation, the water-saturated silicon oil (KF-96-100CS; Shin Etsu Chemical) was layered, and the pollen tubes were subjected to the guidance assay by micromanipulation. Under observation using an inverted microscope (Observer Z1; Zeiss), a single ovule was picked up using a glass needle produced with a glass needle puller (MCF-100; Nepagene) and placed in front of the pollen tubes using a manipulator (MTK-1SH; Narishige).

**Aniline blue staining of in vitro and semi-in vitro pollen-tube growth**

To observe the pollen tubes in vitro, pollen grains were germinated on the growth medium [modified Nitsch's medium containing 13% polyethylene glycol 4000 (w/v), and 1% sucrose (w/v)] containing AMOR (TCI) or multivalent AMORs with 1.5% ultla-low gelling agarose (agarose type IX-A; Sigma) and incubated at dark at 28 °C for 16 h. To observe the pollen tubes semi-in vitro, the cut end of 15-mm-long hand-pollinated style was placed on the growth medium containing AMOR (TCI) or multivalent AMORs with 1.5% ultla-low gelling agarose and incubated at dark at 28 °C for 16 h. After incubation, callose plugs in pollen tubes were labelled using 0.1% aniline blue dye (in 0.1 M $K_2HPO_4$) and observed by Observer Z1 (Zeiss).
